# Capsular polysaccharide inhibits vaccine-induced O-antigen antibody binding and function across both classical and hypervirulent K2:O1 strains of *Klebsiella pneumoniae*

**DOI:** 10.1101/2022.11.01.514596

**Authors:** Paeton L. Wantuch, Cory J. Knoot, Lloyd S. Robinson, Evgeny Vinogradov, Nichollas E. Scott, Christian M. Harding, David A. Rosen

**Affiliations:** Department of Pediatrics, Division of Infectious Diseases, Washington University School of Medicine, St. Louis, MO 63110, USA; Omniose, St. Louis, MO, 63110, USA; National Research Council Canada, Human Health Therapeutics Centre, Ottawa, ON K1A 0R6, Canada; Department of Microbiology and Immunology, The Peter Doherty Institute for Infection and Immunity, University of Melbourne, Parkville, VIC 3010, Australia; Department of Molecular Microbiology, Washington University School of Medicine, St. Louis, MO 63110. USA

**Keywords:** *Klebsiella pneumoniae*, vaccine, capsular polysaccharide, O-antigen polysaccharide, bioconjugation

## Abstract

*Klebsiella pneumoniae* presents as two circulating pathotypes: classical *K. pneumoniae* (c*Kp*) and hypervirulent *K. pneumoniae* (hv*Kp*). Classical isolates are considered urgent threats due to their antibiotic resistance profiles, while hv*Kp* isolates have historically been antibiotic susceptible. Recently, however, increased rates of antibiotic resistance have been observed in both hv*Kp* and c*Kp*, further underscoring the need for preventive and effective immunotherapies. Two distinct surface polysaccharides have gained traction as vaccine candidates against *K. pneumoniae*: capsular polysaccharide and the O-antigen of lipopolysaccharide. While both targets have practical advantages and disadvantages, it remains unclear which of these antigens included in a vaccine would provide superior protection against matched *K. pneumoniae* strains. Here, we report the production of two bioconjugate vaccines, one targeting the K2 capsular serotype and the other targeting the O1 O-antigen. Using murine models, we investigated whether these vaccines induced specific antibody responses that recognize K2:O1 *K. pneumoniae* strains. While each vaccine was immunogenic in mice, both c*Kp* and hv*Kp* strains exhibited decreased O-antibody binding in the presence of capsule. Further, O1 antibodies demonstrated decreased killing in serum bactericidal assays with encapsulated strains, suggesting that the presence of *K. pneumoniae* capsule blocks O1-antibody binding and function. Finally, the K2 vaccine outperformed the O1 vaccine against both c*Kp* and hv*Kp* in two different murine infection models. These data suggest that capsule-based vaccines may be superior to O-antigen vaccines for targeting hv*Kp* and some c*Kp* strains, due to capsule blocking the O-antigen.

**Significance Statement:** Currently there are no licensed vaccines targeting *K. pneumoniae*, but several are in development. Two prominent *K. pneumoniae* surface polysaccharides (capsule and O-antigen) represent attractive vaccine targets; however, the relative efficacy of these potential vaccines against *K. pneumoniae* strains has not been directly compared. To inform future vaccine development, we evaluate two bioconjugate vaccines (targeting either capsule or O-antigen) demonstrating that each are immunogenic in murine models. However, we find that *K. pneumoniae* capsule largely inhibits recognition by antibodies raised against O-antigen. Further, we demonstrate that a capsule-based vaccine outperforms an O-antigen vaccine against both c*Kp* and hv*Kp* in murine models of pneumonia and bacteremia, suggesting that capsule-based vaccines offer superior protection from some *K. pneumoniae* infections.

## Introduction

*Klebsiella pneumoniae* is an encapsulated, Gram-negative, opportunistic pathogen that causes an array of human infections. It has recently been recognized as the most common contributory pathogen in deaths from infectious causes in children under 5 years of age (*1*). Two distinct pathotypes of *K. pneumoniae* are currently circulating globally: classical *K. pneumoniae* (c*Kp*) and hypervirulent *K. pneumoniae* (hv*Kp*) (*2*). c*Kp* is known for causing healthcare-associated infection and is notorious for the capacity to express a variety of antibiotic resistance determinants such as extended-spectrum beta-lactamases (ESBLs) or *K. pneumoniae* carbapenemases (KPCs) (*3–5*). In contrast, hv*Kp* typically manifests as community-acquired infection in otherwise healthy hosts. Historically, hv*Kp* strains were sensitive to most antibiotics; however, genetic exchanges between hv*Kp* and c*Kp* have recently produced hv*Kp* isolates resistant to most, if not all, antibiotics (*6, 7*).

Antibiotic-sparing therapies are desperately needed to prevent and combat *K. pneumoniae* infections, but there is not currently a licensed vaccine targeting *K. pneumoniae*. Several recent studies have proposed putative vaccine candidates, most notably polysaccharides such as capsule or O-antigen, for inclusion into polysaccharide-protein conjugate vaccines (*8–10*). O-antigen remains a leading candidate for incorporation into a vaccine, and at least one tetravalent O-antigen-based bioconjugate vaccine is currently in Phase I trials (*11*). However, given the lack of head-to-head studies, the most effective *K. pneumoniae* polysaccharide antigen has yet to be definitively identified.

Capsular polysaccharide is a critical *K. pneumoniae* virulence factor whose structure can vary significantly among strains. Of note, hv*Kp* isolates typically produce extremely high levels of capsular polysaccharide on their surfaces and often exhibit a hypermucoviscous appearance (*12*). This exuberant production of capsule is directly linked to impairment of complement-mediated killing (*13*) and phagocytosis (*14*). Over 100 distinct capsule serotypes have been described for *K. pneumoniae* to date (*15, 16*). However, one serotype of note, the K2 capsular type, is associated with the majority of hv*Kp* infections as well as ~8% of c*Kp* infections (*12, 17*).

The other major surface polysaccharide of *K. pneumoniae*, O-antigen, is incorporated into the terminus of lipopolysaccharide (LPS) which consists of the lipid A molecule, a core saccharide, and the terminal repeating O-antigen polysaccharide (*18*). O-antigen is characterized into different serogroups based on distinct structures and antigenic properties. In contrast to the capsular polysaccharide, *K. pneumoniae* has only eleven characterized LPS serogroups, with over 80% of all isolates expressing just four serogroups (*17, 19, 20*). One of these four is the O1 O-antigen.

Little is known about relative exposure or direct interactions between the capsular and O-antigen polysaccharides on the bacterial cell surface. However, one investigation demonstrated that O1-specific antibodies had differential effects on surface properties of *K. pneumoniae* depending on whether the capsular type was K1 or K2 (*21*), suggesting that the structure of capsular polysaccharide could affect interaction with O-antigen antibodies. Additional studies have shown that monoclonal antibodies to galactan II (a component of the O1 O-antigen) were able to bind O1 *K. pneumoniae* strains lacking capsule but did not bind, or bound significantly less, to strains expressing K2, K7, or K21 capsules (*22*). Together, these studies suggest possible interactions between capsule, O-antigen, and their respective targeting antibodies, with important implications for vaccine efficacy.

Here we investigated the relative efficacy of *K. pneumoniae* polysaccharide-protein conjugate vaccines to determine which polysaccharide is a superior vaccine target. We developed glycoconjugate vaccines targeting the K2 capsule and O1 O-antigen using bioconjugation, an enzymatic approach to generating polysaccharide-protein conjugates *in vivo* using *E. coli* as a host (*23*). Here, we present our newly designed glycoengineering approach used in constructing these vaccines and demonstrate that while both vaccines produce robust serotype-specific antibody responses, the K2 vaccine outperforms the O-antigen vaccine in multiple murine models of *K. pneumoniae* infection. ELISA and serum bactericidal data indicate that *K. pneumoniae* capsule itself impairs both the binding and function of O-antigen antibodies. These data will inform future vaccine formulations to target the urgent global threat of *K. pneumoniae*.

## Results

### Heterologous expression of K2 and O1 polysaccharides in *E. coli*

The production of polysaccharide-protein conjugates via bioconjugation requires glycoengineered *E. coli* to express a lipid-linked polysaccharide precursor that is specific to the pathogen of interest. This polysaccharide is then enzymatically transferred by an oligosaccharyltransferase (termed an OTase or conjugating enzyme) to proteins engineered to contain a sequon (an amino acid sequence recognized specifically by the OTase).

To heterologously express the K2 capsular polysaccharide in glycoengineered *E. coli*, we previously reported that co-expression of the *K. pneumoniae* RmpA transcriptional activator was required for K2 capsule expression when the K2 gene cluster (*wcuF-ugd*) was expressed as a single locus (**Fig. 1A**) (*8*). Here, the requirement for RmpA co-expression was overcome by cloning the K2 gene cluster in discrete sections from *wcuF* through *wzx* and *wcaJ* through *ugd* into two separate IPTG-inducible expression vectors (**Fig. 1B**). This cloning strategy was implemented to remove two open reading frames located amid the K2 gene cluster that encode for a hypothetical protein and putative acetyltransferase, as previous NMR analysis of the K2 capsule from both *K. pneumoniae* ATCC 43816 and glycoengineered expressing the K2 polysaccharide *E. coli* did not show acetylation of the K2 repeat unit (*8*). Silver staining and western blot analysis (using historic antisera derived from mice immunized with our previous K2-EPA bioconjugate (*8*)) of LPS extracted from glycoengineered *E. coli* expressing the decoupled K2 locus were found to express significantly more K2 polysaccharide than glycoengineered *E. coli* expressing the K2 gene cluster from a single locus, with or without *rmpA* expression (**Fig. 1C**). Importantly, the K2 polysaccharide produced by glycoengineered *E. coli* containing the decoupled K2 gene cluster was identical to the published structure of the native K2 capsular polysaccharide (*8*) (***SI Appendix***, **Fig. S1**).

**Figure 1.**
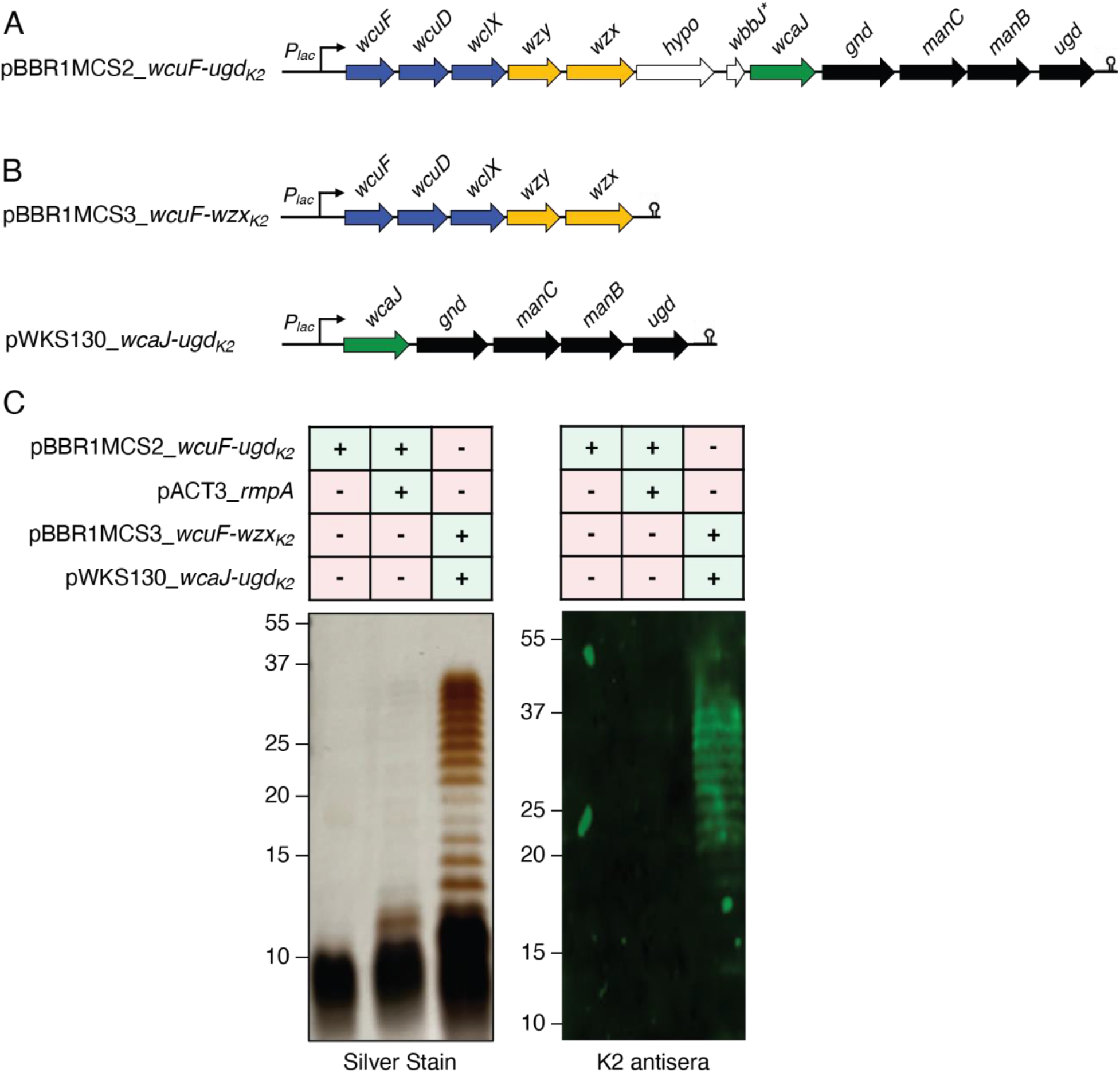
Decoupling the K2 gene cluster over two expression vectors obviates the requirement of RmpA for K2 polysaccharide expression in *E. coli*. A) The K2 gene cluster from *wcuF* to *ugd* was previously cloned as a single locus into the expression vector pBBR1MCS2. The asterisk indicates a putative gene designation of *wbbJ* for the open reading frame immediately upstream of *wcaJ*. (B) A new cloning strategy separated the K2 cluster over two expression vectors and simultaneously removed two open reading frames encoding for a hypothetical protein and the putative acetyltransferase (*wbbJ*). (C) Silver staining and western blot analysis of LPS extracted from glycoengineered *E. coli* expressing the K2 locus as a single locus (with and without RmpA) or the K2 locus separated over two expression vectors.

The *K. pneumoniae* O1v1 O-antigen consists of two types of polysaccharides: an O1 polymer containing the disaccharide repeat unit [→3)-β-D-Gal*p*-(1→3)-α-D-Gal*p*-(1→] (D-galactan II), attached at the non-reducing terminus of a pre-assembled O2a O-antigen polymer with the disaccharide repeat unit [→3)-β-D-Gal*f*-(1→3)-α-D-Gal*p*-(1→] (D-galactan I) (*20, 24*). Like all *K. pneumoniae* O-antigens, O1 is linked to undecaprenol pyrophosphate via a single *N*-acetylglucosamine (GlcNAc) monosaccharide at the polymer reducing end (*25*). Expression of five genes, across two separate genomic loci, is sufficient to produce the *K. pneumoniae* O1 O-antigen in *E. coli (24): wzm, wbbM, glf, wbbO* (located in one locus) and *wbbY* in the other. We cloned the *wbbY* gene into pACT3 (pACT3-*wbbY*) and the remaining four genes into pBBR1MCS2 (pBBR1MCS-O2a) under IPTG-inducible promoters, then expressed these in an *E. coli* K12 derivative. To validate the structure of the glycoengineered O1 polysaccharide produced by *E. coli*, we extracted the O1-containing lipopolysaccharide and analyzed it by nuclear magnetic resonance to determine the structure and linkage. As shown in the overlap of the ^1^H-^13^C HSQC spectra of the *K. pneumoniae* O1 and D-galactan I found in ***SI Appendix*, Fig. S2**, the O1 polysaccharide extracted from glycoengineered *E. coli* matched with the published structure of the native O1 polysaccharide (*24*).

### K2- and O1-bioconjugate vaccine production and characterization

Previously, we demonstrated that a genetically inactivated mutant (ΔE553) of exotoxin A from *Pseudomonas aeruginosa* (EPA) served as an immunogenic carrier protein for the PglS bioconjugation system (*8*) and also described variants of EPA that contain the PglS sequon at various locations in the EPA coding sequence (*23, 26*). For the present work, we integrated two 23-amino acid sequons from the native PglS substrate ComP between EPA residues A489/R490 and E548/G549 (**Fig. 2A**). Prior studies showed that these internal sequons were readily glycosylated by PglS (*26*). To produce O1-EPA and K2-EPA bioconjugates, we paired the plasmids for expressing the O1 polysaccharide or the K2 polysaccharide with a plasmid expressing the EPA protein (modified with two sequons) and the PglS OTase in CLM24 or SDB1 *E. coli* strains (*27*), respectively. Bioconjugates were purified from periplasmic extracts using immobilized metal affinity, anion-exchange, and size-exclusion chromatography. Western blotting of the resulting purified glycoproteins revealed little or no contamination with unglycosylated EPA (**Fig. 2B – D**). The purified O1-EPA bioconjugates reacted with a previously developed mouse monoclonal antibody specific to O1 polysaccharide (*20*) (**Fig. 2C**). Neither antisera or monoclonal antibodies specific to the K2 polysaccharide alone are commercially available, and therefore K2 western blots were unable to be performed.

**Figure 2.**
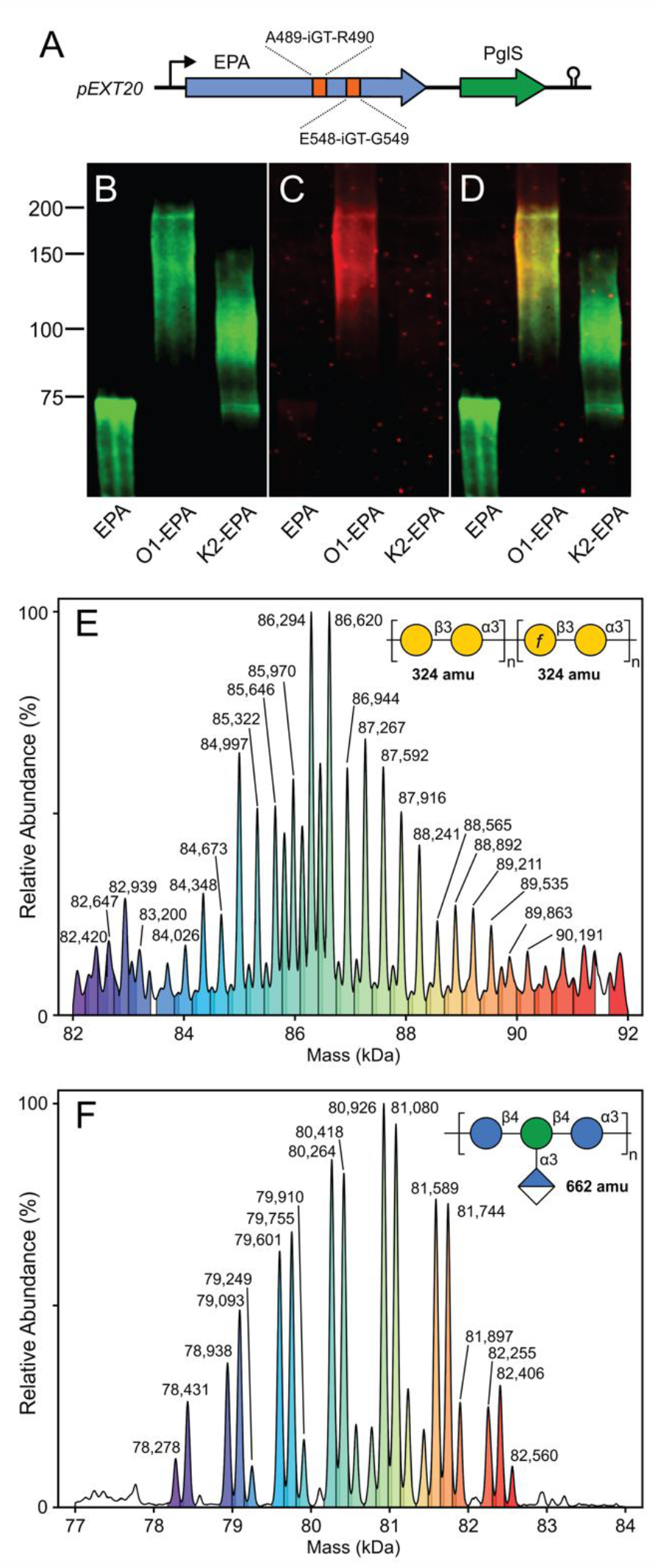
Characterization of *K. pneumoniae* O1-EPA and K2-EPA bioconjugates. A) Diagram of genes encoding EPA and PglS (colored arrows) on the pEXT20-derived expression plasmid. EPA was engineered to contain two internal sequons (orange blocks). Gene expression is driven from an IPTG-inducible promoter (black arrow) with a downstream *rrnB T2* terminator (black hairpin). (B) Western blot of purified EPA and *K. pneumoniae* bioconjugates, anti-EPA channel. (C) Western blot, anti-O1 channel. (D) Merged channels from panels B and C. (E) MS1 spectrum of intact, purified O1-EPA bioconjugate. A diagram of the O1 repeat unit is inset in the top right of the panel. (F) MS1 spectrum of intact, purified K2-EPA bioconjugate. A diagram of the K2 repeat unit is inset in the top right of the panel.

Intact protein mass spectrometry was used to further characterize the purified O1-EPA and K2-EPA bioconjugates (**Fig. 2E-F**). The covalent attachment of glycans containing different numbers of repeating K2 or O1 units is readily observable in the MS1 spectrum of the bioconjugates, enabling relative polysaccharide to protein ratios to be characterized. Typically, bioconjugates exhibit a gaussian-like distribution of repeating sugar units, with the mean number of repeat units varying between polysaccharides. Intact MS1 spectrum of the O1-EPA (**Fig. 1E**) displayed a glycoprofile with major peaks separated by ~324 amu, the theoretical mass of the constituent O1 disaccharide repeating unit. Both galactan I (O2a portion) and galactan II (the O1-specific addition at the non-reducing terminus) consist of repeating galactose disaccharide units and therefore have equal mass, 324 amu. Using a calculated mass of 71,509 amu for EPA and accounting for the priming GlcNAc monosaccharide at the reducing end, we estimate that most of the O1 glycans attached to EPA contain between 39 to 55 disaccharide repeating units corresponding to peaks with masses 84,348 amu to 89,535 amu. Intact MS1 spectrum of K2-EPA (**Fig. 2F**) also revealed a gaussian-like glycoform profile with major peaks separated by ~662 amu, equal to the calculated mass of the K2 tetrasaccharide repeat unit. We estimated that most K2 glycans contained between 10 to 16 repeat units corresponding to peaks with masses 78,278 amu to 82,255 amu. Using the ion intensities and areas under each curve as well as the observed masses of each glycoform (carrier protein mass + polysaccharide mass), we calculated the contribution of the polysaccharide portion to each glycoform’s total ion intensity. Using this method, polysaccharide-to-protein ratios were found to be 0.256 and 0.144 for the O1-EPA and K2-EPA bioconjugates, respectively.

### Capsule of hv*Kp* inhibits binding of O-antigen IgG

To begin comparing responses to capsule and O-antigen bioconjugate vaccines, we immunized BALB/c mice in three groups: control immunization (unglycosylated EPA protein), K2-EPA bioconjugate, or O1-EPA bioconjugate. The K2-EPA and O1-EPA bioconjugate vaccines were each formulated to contain 1 μg of polysaccharide. All vaccines were formulated with Alhydrogel (2% aluminum hydroxide gel) as an adjuvant at a 1:9 ratio. Mice were immunized on days 0, 14, and 28. Serum samples were collected prior to each immunization and on day 42 and were used to determine levels of polysaccharide-specific IgG via ELISA on plates coated with *K. pneumoniae* 43816, a K2:O1 isolate. The 43816 strain has been extensively used as a model strain for studying hv*Kp* virulence in murine infections (*8*).

In mice immunized with the K2-EPA bioconjugate, we observed robust levels of K2-specific IgG that significantly increased over the course of immunization, compared to pre-immune sera (**Fig. 3A**). As expected, we observed no significant 43816-specific IgG response in control EPA-immunized mice. Interestingly, we detected no significant IgG binding to 43816 by the sera of O1-EPA-vaccinated mice. As some hv*Kp* are known to be hypermucoviscous, we hypothesized that capsule may be blocking the O-antigen. To test this hypothesis, or alternatively, to determine if O1-EPA failed to induce O1-specific antibodies, we derived a capsule-deficient mutant from the parent 43816 strain using a modified Red recombinase method (*28*). The 43816Δ*cps* mutant lacks the entire K2 capsular locus, rendering it less mucoviscous and unable to produce capsule as demonstrated by uronic acid quantification (***SI Appendix*, Fig. S3**). Using the sera from vaccinated mice described above, ELISAs using plates coated with 43816Δ*cps* now demonstrated robust IgG binding by the sera of O1-EPA-immunized mice that increased over the course of immunization (**Fig. 3B**). As expected, we did not observe any specific IgG recognizing 43816Δ*cps* from the sera of EPA- or K2-EPA-immunized mice. Importantly, repeating the ELISA using a mutant unable to synthesize the O1 antigen (43816Δ*wecA*) as the coating agent did not demonstrate any binding by sera from O1-EPA immunized mice (**Fig. 3C**). To confirm the presence of the expected polysaccharide antigens in each of the mutants and the specificity of the immune sera, western blots were performed on whole-cell lysates of 43816, 43816Δ*cps*, and 43816Δ*wecA* probing with K2-EPA mouse sera (**Fig. 3D**) or O1-EPA mouse sera (**Fig. 3E**). Corroborating the ELISA results, we observed strong K2-EPA seroreactivity to 43816 and 43816Δ*wecA*, as well as strong O1-EPA seroreactivity to 43816 and 43816Δ*cps*. Taken together, these data indicate that the O1-EPA bioconjugate elicited robust O1-specific IgG responses, but that O1-specific IgG was unable to bind to 43816 due to K2 capsule blocking O1-specific IgGs from binding their antigenic target.

**Figure 3.**
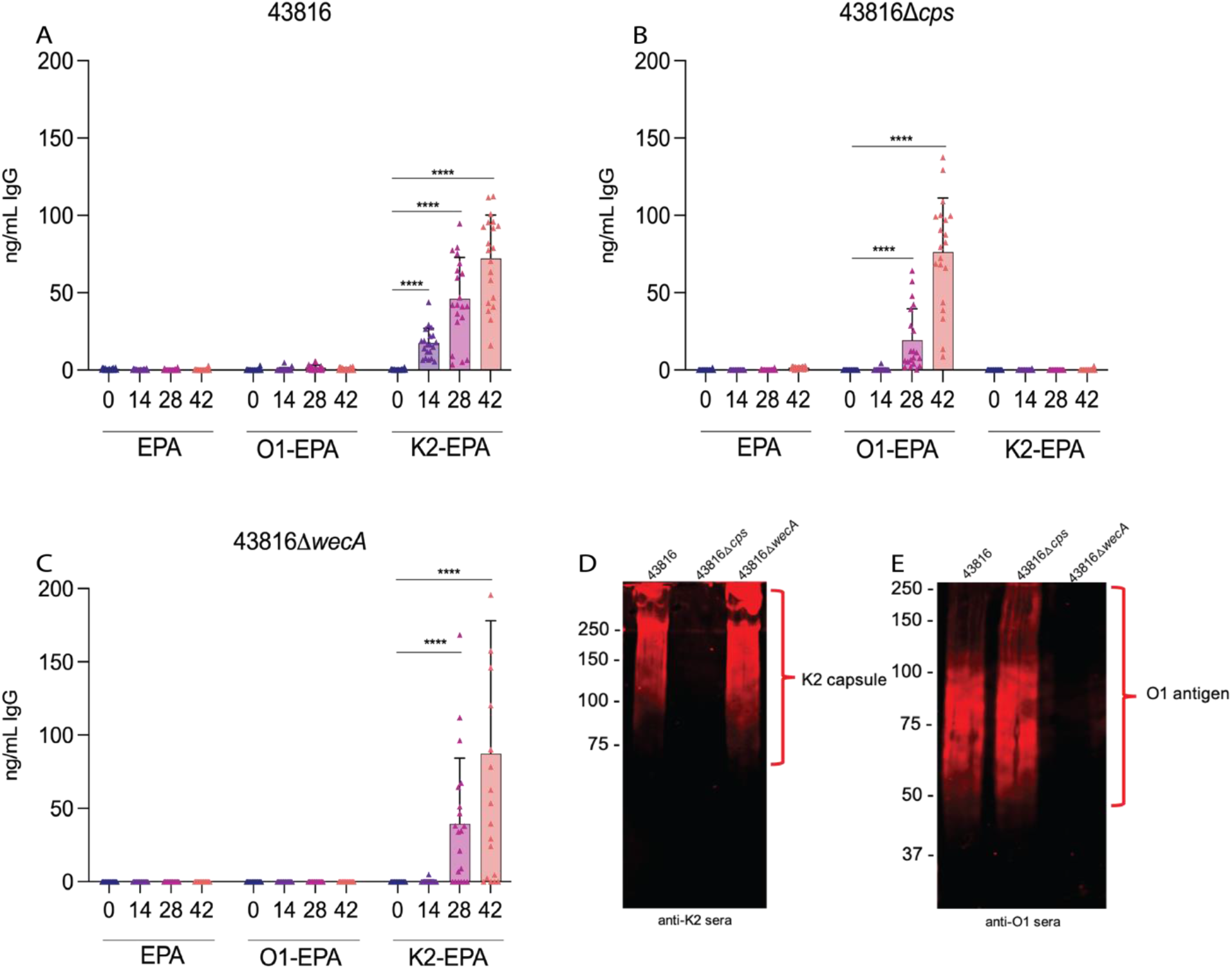
Detection of IgG generated by O1-bioconjugate and K2-bioconjugate vaccines. Carrier protein (EPA), O1- or K2-specific IgG kinetics over the course of the immunization series as measured by ELISA against whole bacteria of strains (A) 43816, (B) 43816Δ*cps*, or (C) 43816Δ*wecA*. Western blot analyses probing for (D) K2 glycan using pooled day 42 sera from K2-vaccinated mice or (E) O1 glycan using pooled day 42 sera from O1-vaccinated mice. Statistical analyses were performed via Mann-Whitney U test. **** p<0.0001. Error bars represent standard deviations.

### Capsule of c*Kp* impairs binding of O-antigen IgG

To determine if the phenomenon of capsule shielding of O-antigen was specific to the more heavily encapsulated hv*Kp* strains, we investigated antisera binding to a c*Kp* isolate with the same capsule and O-antigen types (K2:O1). KR174 is a c*Kp* that was isolated from the lower respiratory tract of a patient with c*Kp* pneumonia and encodes an ESBL (*29*). It produces less capsule and displays decreased hypermucoviscosity relative to hv*Kp* 43816 (***SI Appendix***, **Fig. S3**). We performed ELISA with the sera described above, using plates coated with c*Kp* KR174. As expected, sera from K2-EPA-immunized mice exhibited a robust IgG response that significantly increased over the course of immunization, while no reactivity was observed in sera from the EPA-immunized group (**Fig. 4A**). Interestingly, sera from mice immunized with the O1-EPA exhibited minor IgG responses to KR174 that appeared over the course of immunization (**Fig. 4A**); however, these levels were significantly lower than those observed with K2-EPA sera (p<0.0001). As before, to test if capsule could be blocking O-antigen in this classical strain of *K. pneumoniae*, we deleted the capsular locus in KR174 and tested IgG binding. Indeed, we observed no significant IgG in the sera of mice vaccinated with EPA or K2-EPA when ELISA was performed using KR174Δ*cps* as the coating agent (**Fig. 4B**); however, almost 10-fold higher O1-EPA IgG concentrations were detected by ELISA against KR174Δ*cps* when compared to wild-type KR174. Repeating the ELISA using KR174Δ*wecA* as the coating agent resulted in the loss of detectable IgG in sera of O1-EPA immunized mice (**Fig. 4C**). Western blots revealed that K2-EPA sera reacted strongly against whole-cell lysates of KR174 and KR174Δ*wecA* but not KR174Δ*cps* (**Fig. 4D**), and that O1-EPA sera reacted strongly reacted against whole-cell lysates of KR174 and KR174Δ*cps*, but not KR174Δ*wecA* (**Fig. 4E**). These data suggest that, as in hv*Kp*, capsule in this c*Kp* strain is largely blocking O-antigen recognition by O1-specific antibodies.

**Figure 4.**
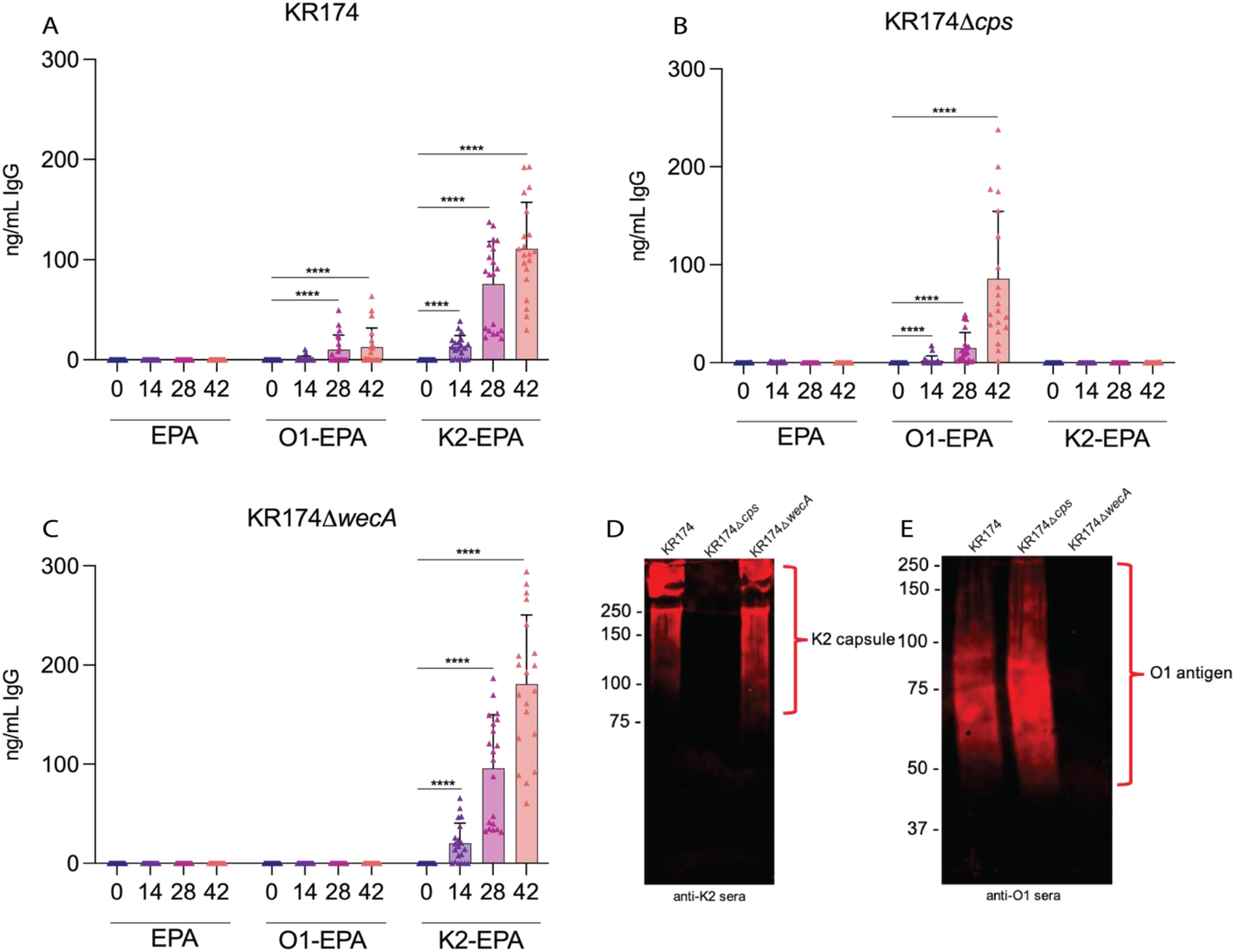
Detection of IgG generated by O1-bioconjugate and K2-bioconjugate vaccines. Carrier protein (EPA), O1 or K2-specific IgG kinetics over the course of the immunization series as measured by ELISA against whole bacteria of strains (A) KR174, (B) KR174Δ*cps*, or (C) KR174Δ*wecA* Western blot analyses probing for (D) K2 glycan using pooled day 42 sera from K2-vaccinated mice or (E) O1 glycan using pooled day 42 sera from O1-vaccinated mice. Statistical analyses were performed via Mann-Whitney U test. **** p<0.0001. Error bars represent standard deviation.

### Capsule inhibits O1-specific antibody function

As differences in anti-K2 and anti-O1 antibody binding were observed by ELISA, we next sought to determine if these antisera demonstrated functional differences against *K. pneumoniae*. Serum bactericidal assays (SBAs) with wild-type strains 43816 or KR174, and their isogenic mutants, were used to quantify the ability of vaccine-elicited antibodies to evoke complement-mediated killing (**Fig. 5**). We observed very robust and significant killing (~75%) of 43816 by K2-EPA-derived antibodies, with no significant killing from antibodies generated by control EPA vaccination (**Fig. 5A**). Surprisingly, O1-EPA-derived antibodies mediated modest (~25%) bactericidal activity, but was significantly less killing compared to K2-EPA sera (**Fig. 5A**). To determine if capsule was inhibiting more robust killing by anti-O1 antibodies, we repeated the SBAs utilizing 43816Δ*cps*. As expected, we observed no significant killing mediated by EPA sera or anti-K2 sera (**Fig. 5B**). Loss of capsule, however, resulted in greater O1 mediated killing (~55%) of 43816Δ*cps* (**Fig. 5B**) compared to that observed in 43816. Further, in SBAs using 43816Δ*wecA* (**Fig. 5C**) we observed a loss of bacterial killing mediated by O1-EPA sera, re-confirming the O1 antigen specificity and functionality of these O1 antibodies.

**Figure 5.**
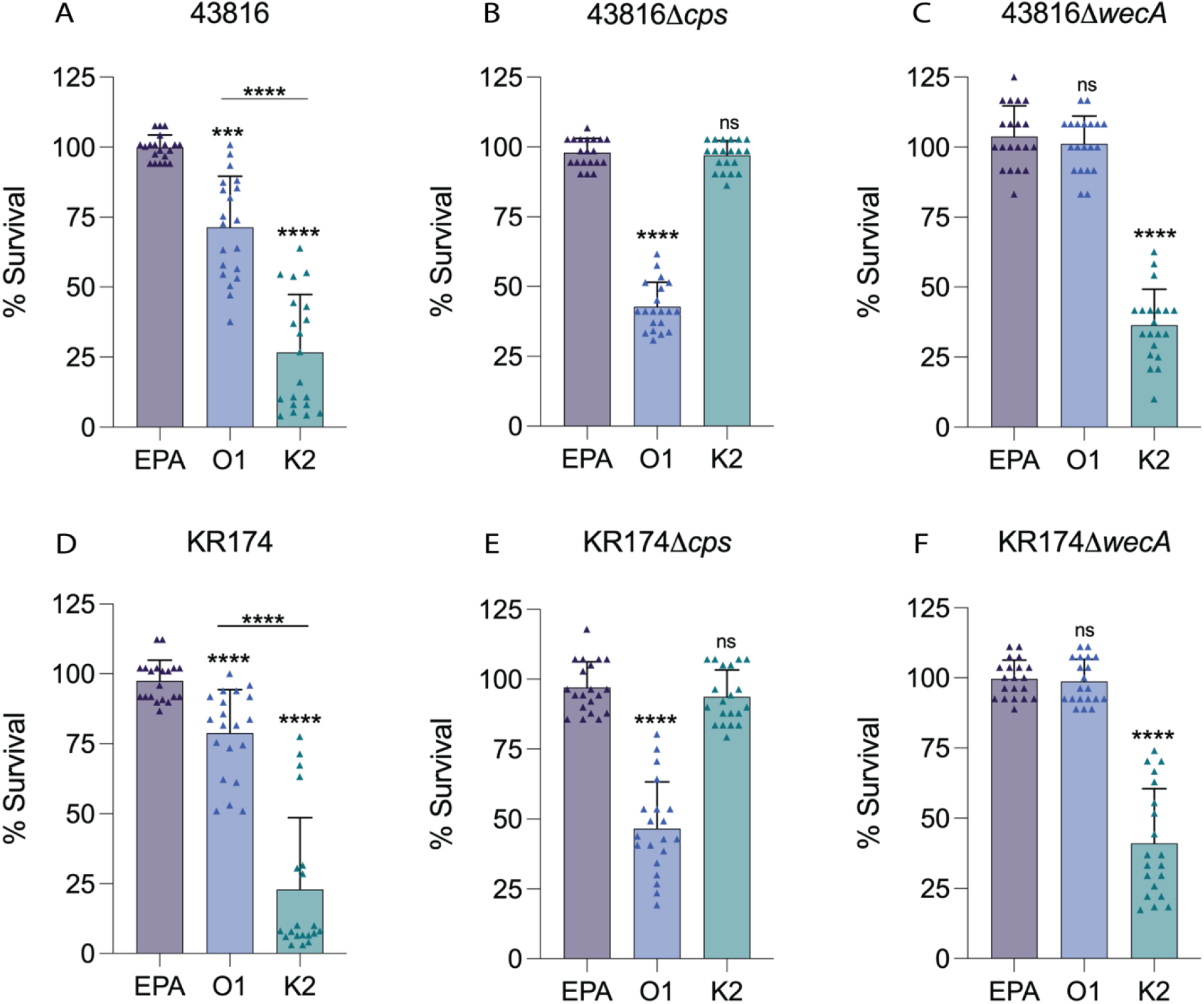
Serum bactericidal assays with bioconjugate-vaccinated mouse sera. Serum bactericidal activities of day 42 sera from mice immunized with EPA carrier protein, O1-EPA, or K2-EPA bioconjugates were measured against (A) 43816, (B) 43816Δ*cps*, (C) 43816Δ*wecA*, (D) KR174, (E) KR174Δ*cps*, or (F) KR174Δ*wecA*. Each data point represents a single mouse. Statistical analyses were performed via Mann-Whitney U tests in comparison to EPA survival, unless otherwise indicated. *** p<0.001; **** p<0.0001; ns, not significant. Error bars represent standard deviation.

We next performed SBAs with KR174 to determine if these immune sera would perform similarly against a classical isolate of *K. pneumoniae*. As expected, we observed no significant killing by control EPA-immunized sera but robust (~75%) killing with anti-K2 sera (**Fig. 5D**). As with 43816, we observed modest (~20%) killing of KR174 mediated by anti-O1 antisera, significantly less than with anti-K2 antisera (**Fig. 5D**). Using KR174Δ*cps*, we detected greater bacterial killing mediated by the anti-O1 antisera (~55%) compared to that observed in KR174, and none with the anti-K2 antisera (**Fig. 5E**). Finally, O1-EPA antisera did not enhance complement-mediated killing of KR174Δ*wecA*, demonstrating their specificity for the O1 O-antigen (**Fig. 5F**). Together these data support a model in which capsule is indeed blocking O-antigen to varying degrees in both hv*Kp* and c*Kp* strains, resulting in diminished binding and function of O-antigen-specific antibodies, and that antibodies targeting the capsule exert superior binding and killing of both c*Kp* and hv*Kp*.

### K2-bioconjugate vaccine provides superior protection against hv*Kp* pneumonia

Little is known about correlates of protective immunity to either classical or hypervirulent *K. pneumoniae* infections. Therefore, to determine if capsule or O-antigen-based bioconjugate vaccines confer protection from pneumonia, we evaluated their efficacy in a murine pulmonary challenge model using either hv*Kp* 43816 or c*Kp* KR174 at inocula close to their LD_90_ values. 43816 has an LD_90_ in this model of ~1,500 colony-forming units (CFU) (*8, 30*), and we observed separately that KR174 doses of ~10^9^ CFU in BALB/c mice killed the vast majority of mice. Three groups of mice were immunized with the carrier protein alone (EPA), K2-EPA, or O1-EPA bioconjugate vaccines. Mice were immunized on days 0, 14, and 28, then challenged on day 42 with a lethal dose of either 43816 or KR174 via oropharyngeal aspiration and monitored for survival and weight loss for two weeks (**Fig. 6A**). Among K2-EPA immunized mice, 90% survived 43816 pulmonary infection compared to just 15% of EPA-immunized mice and 5% of O1-EPA-immunized mice (p<0.0001) (**Fig. 6B**). Meanwhile, upon challenge with the c*Kp* strain KR174, both O1-EPA and K2-EPA vaccinated mice showed more modest but significant protection relative to control mice (**Fig. 6C**). Specifically, while 90% of EPA-immunized mice succumbed to infection with KR174, we observed 25% protection in O1-EPA immunized mice (p= 0.0208) and 35% protection in K2-EPA immunized mice (p=0.0008). There was no significant difference in protection between O1-EPA and K2-EPA immunized mice (p=0.2164; **Fig. 6C**).

**Figure 6.**
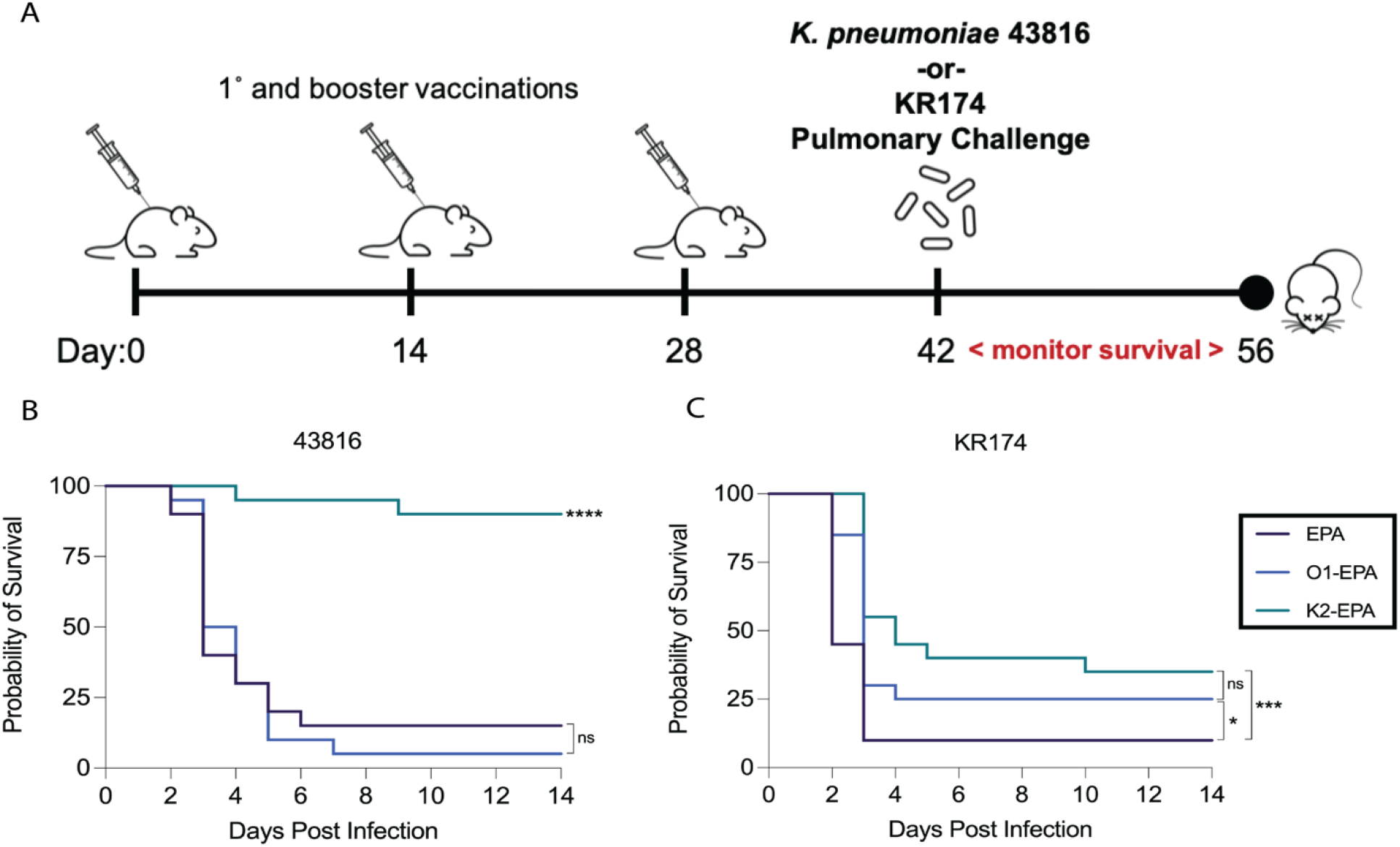
Survival of bioconjugate-vaccinated mice after lethal pulmonary challenge with hv*Kp* or c*Kp*. (A) Mice were vaccinated with either the carrier protein alone (EPA) or the O1-EPA or K2-EPA bioconjugates on days 0,14, and 28, followed by oropharyngeally aspirated with *K. pneumoniae* 43816 or KR174 and monitored for survival for 14 days. (B) Mice were infected with ~1500 CFU for 43816 or (C) ~10^9^ CFU for KR174. Each group contains n=20 mice combined over two independent experiments. Statistical analyses were performed via log-rank (Mantel-Cox) tests comparing against EPA group, unless otherwise indicated. **** p<0.0001; ***p<0.001; * p<0.05; ns, not significant.

### K2-bioconjugate vaccine is superior in a *K. pneumoniae* bacteremia model

To ensure that our vaccine efficacy findings were not specific to a pulmonary challenge model, we employed an additional murine model of bacteremia and dissemination. Groups of BALB/c mice were immunized with the same regimen as described above (**Fig. 7A**). Infection was initiated via intraperitoneal injection, which results in universal bacteremia (***SI Appendix*, Fig. S4**). As the lethal i.p. dose with these strains was unknown, we determined doses of ~2,000 CFU for 43816 and ~10^8^ CFU for KR174 after trial doses at these inocula resulted in almost complete mortality of mice. Mice immunized with EPA alone or the O1-EPA bioconjugate experienced 100% mortality when challenged i.p. with 43816; in contrast, 100% of mice immunized with K2-EPA survived (p<0.0001) (**Fig. 7B**).

**Figure 7.**
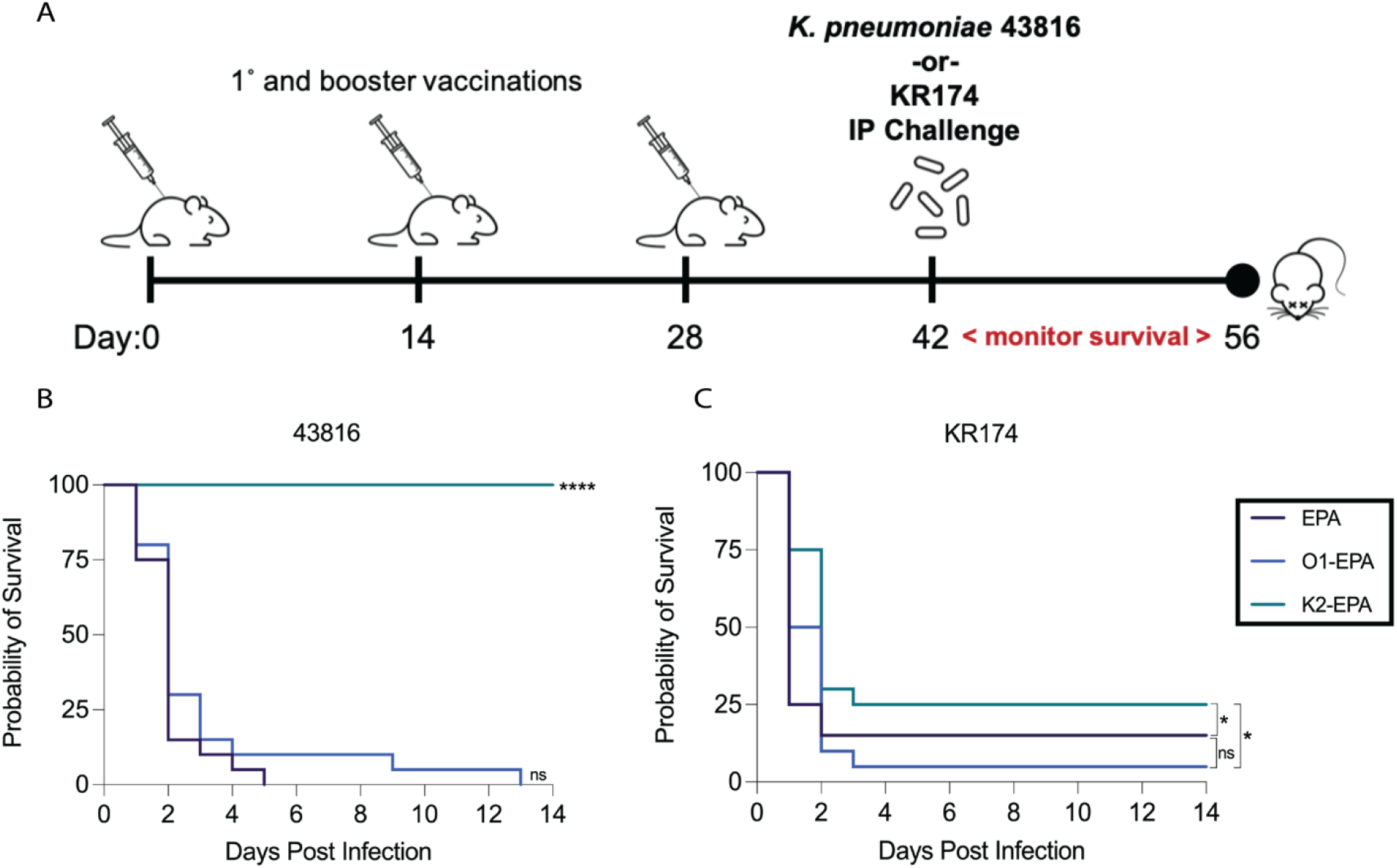
Survival of bioconjugate-vaccinated mice after lethal bacteremia challenge with hv*Kp* or c*Kp*. (A) Mice were vaccinated with either the carrier protein alone (EPA) or the O1-EPA or K2-EPA bioconjugates on days 0, 14, and 28, followed by intraperitoneal injection with *K. pneumoniae* 43816 or KR174 and monitored for survival for 14 days. Mice were infected with (B) ~2000 CFU of 43816 or (C) ~10^8^ CFU of KR174. Each group contains n=20 mice combined over two independent experiments. Statistical analyses were performed via log-rank (Mantel-Cox) tests comparing against EPA group, unless otherwise indicated. **** p<0.0001; * p<0.05; ns, not significant.

As was seen in the pneumonia model, experiments with c*Kp* strain KR174 yielded lower survival overall (**Fig. 7C**). The K2-EPA bioconjugate did demonstrate modest, statistically significant protection compared to EPA-immunized mice (p=0.0435) and O1-EPA-immunized mice (p=0.0377). Together, these data support the use of a matched, capsule-based bioconjugate vaccine over an O-antigen-based vaccine to protect mice from bacteremia with c*Kp* or hv*Kp*.

## Discussion

*K. pneumoniae* is a versatile bacterium that causes many human infections including pneumonia, urinary tract infection and sepsis. Treatment of this pathogen has become increasingly more difficult as antibiotic resistance continues to rise. *K. pneumoniae* is the most prevalent of the carbapenem-resistant Enterobacterales (CRE) (*31*), and the CDC has designated CRE one of five urgent-level threats in the United States. It has become evident that two distinct pathotypes of *K. pneumoniae* are circulating, c*Kp* and hv*Kp*. c*Kp* isolates are often associated with nosocomial infections, especially among immunocompromised individuals; and all nosocomial *K. pneumoniae* infections are on the rise. In fact, *K. pneumoniae* is now the most common cause of nosocomial pneumonia (*32*). In contrast, hv*Kp* isolates cause a range of disseminated, community-acquired infections in otherwise healthy individuals (*33*). Even more alarming is the emergence of hv*Kp* strains resistant to most or all antibiotics (*6*). Continuing with this theme, it was recently determined that *K. pneumoniae* is now the most common contributory pathogen to death in children under five-years-old with infectious causes in low- and middle-income countries (*1*).

Vaccines have the potential to prevent *K. pneumoniae* infections in geographically burdened locations and highly susceptible populations. The two prominent surface polysaccharides of *K. pneumoniae*, capsule and O-antigen, serve as the most attractive candidates for vaccine inclusion. While capsule may be the most prominently exposed and important virulence factor, the ability to theoretically target a larger proportion of *K. pneumoniae* clinical isolates with fewer included O-antigens is appealing. However, to our knowledge, the relative efficacy of matched O and K vaccines has not been studied head-to-head. This is especially important as questions about shielding of O-antigen or interactions between capsule and O-antigen have been raised previously (*21, 22, 34*). Here we produced and characterized two bioconjugate vaccines, one targeting the K2 capsule type and the other the O1 O-antigen of *K. pneumoniae*. We demonstrated that while both vaccines produce robust and functional antibody responses, capsule in both c*Kp* and hv*Kp* largely blocks the binding and functionality of O-antigen antibodies. Further, only the K2 vaccine exerted significant protection in murine pulmonary and bacteremia challenge models against hv*Kp*; furthermore, K2 vaccine elicited greater protection in c*Kp* bacteremia (and trended similarly in the pneumonia model) than did the O1 vaccine. Putting these results in context, vaccines based solely on O-antigen may be less effective against many circulating strains, while an ideal multivalent *K. pneumoniae* vaccine may require a combination of capsule antigens with or without additional O-antigen targets. For example, capsule types such as K1 and K2 (*12, 33*), highly prevalent among hv*Kp* isolates, and KL106 and KL107 (*16, 35, 36*), highly prevalent in antibiotic-resistant clonal groups, would be attractive capsular types to test in combination with common O1 and O2, O3, and O5 antigens in a multivalent vaccine (*37*).

The novel bioconjugation technology used to produce the present *K. pneumoniae* vaccine candidates boasts several benefits. First, glycoconjugate vaccines traditionally are manufactured using chemical conjugation (*38*) which is a multi-step, complex, labor-intensive, and expensive production method (*39*). Bioconjugation, on the other hand, relies on an oligosaccharyltransferase to transfer the polysaccharide of interest to a carrier protein all within the periplasm of a bacterial expression system (*23*). The bioconjugate vaccines used in this work were produced with the conjugating enzyme PglS, which has the unique ability to transfer virtually any polysaccharide, including those containing glucose as the reducing end sugar, to engineered carrier proteins (*23, 40*). This is especially important as over 50% of all *K. pneumoniae* capsular serotypes contain glucose at their reducing ends, including K1 and K2 (*15*). Second, we have improved upon the previously published K2 vaccine (*8*) by shortening the glycosylation site (sequon) that PglS recognizes. This newly identified sequon can be placed internally in the carrier protein (as published here) allowing for *bona fide* glycosylation of the EPA carrier protein. This internal sequon improves upon the bioconjugate characteristics, such as protein stability over time and protection from *in vivo* proteolysis which will be critical for scale-up and production. Finally, bioconjugation is advantageous over traditional methods as it allows for quick adaptation to circulating strains or even strain pressure that might emerge over time following introduction of the vaccine itself. As bioconjugation relies upon plasmids within a glycoengineered expression system, there is intrinsic capacity to rapidly generate new vaccine constructs through a simple plasmid switch.

In these studies, we immunized groups of five-week-old BALB/c female mice with either the carrier protein alone (EPA), the K2-bioconjugate, or the O1-bioconjugate. Utilizing ELISAs against whole bacteria we demonstrated that in hv*Kp* the capsule entirely blocked the binding of any O1 specific antibodies. However, upon loss of capsule via genetic manipulation, we observed a robust level of O1 IgG present. Low levels of O-antigen antibodies were able to be detected via ELISA when the c*Kp* isolate was used as the coating agent, but like the hv*Kp* strain, when capsule was removed the level of O1 antibodies binding the target significantly increased. Further, we determined through serum bactericidal assays that both the K2 and O1 antibodies were functional and able to induce complement-mediated killing of *K. pneumoniae*. Corroborating the ELISA data, SBAs revealed that when capsule was removed, the functionality of O1 antibodies greatly increases. Together, all these data support a model in which capsule is inhibiting O-antigen antibody recognition in both c*Kp* and hv*Kp*. Furthermore, this work, to the best of our knowledge, is the first published using vaccine-induced antibodies in SBA assays against *K. pneumoniae*. This is a significant contribution to the field, as *K. pneumoniae* currently lacks standardized functionality assays that correlate with protective immunity. Such assays are well established for other pathogens such as *Streptococcus pneumoniae* (*41*) and are imperative for testing future immunotherapeutics. We aim to continue development of standardized *K. pneumoniae* functional assays and determine their correlation with protective immunity.

Beyond demonstrating the production and functionality of K2- and O1-specific IgG through vaccination, we also explored the protectiveness of these two bioconjugates. We tested protection not only using two strains of *K. pneumoniae* (one c*Kp* and one hv*Kp*) but also using two infection models: pneumonia and bacteremia. As we had shown that a prior-generation K2 bioconjugate vaccine was effective in preventing death from K2 pneumonia (*8*), this work extends our findings to bacteremia. We observed profound protection attributable to the newer K2 bioconjugate in both pneumonia (90% protection) and bacteremia (100% protection) models against a hv*Kp* isolate. Remarkably, the challenge inocula were over 10-fold the established LD_50_ for this pathogen. In both models, however, the O1 bioconjugate failed to protect mice, suggesting this antigen would be a poor choice for targeting hv*Kp* strains. When mice received pulmonary challenge with a very high inoculum of c*Kp* (10^9^ CFU), we observed modest but significant protection from either the K2 or O1 vaccine. Notably, while not statistically significant, the K2-immunized mice had slightly higher survival. In contrast, only the K2 vaccine conferred significant protection in the c*Kp* bacteremia model. These results suggest that a matched capsule vaccine is preferable for protection against this c*Kp* isolate over an O-antigen vaccine.

Past studies have inferred better efficacy of O-antigen glycoconjugate vaccines than we observe here. A previous glycoconjugate vaccine composed of four *K. pneumoniae* O-antigen serogroups (O1, O2, O3 and O5) was developed (*9*). In their study, the authors demonstrated this vaccine was immunogenic, and passive transfer of vaccine-induced antibodies into mice was protective from systemic *K. pneumoniae* infection (*9*). However, this study differs from our study in key ways that preclude direct comparison. The vaccine-induced antibodies were generated in New Zealand White rabbits and then passively transferred to mice. Furthermore, rabbits were immunized with a four-dose regimen, and with five times the polysaccharide concentration per dose (5 μg) and Freund’s adjuvant (rather than alum), compared to the three-dose regimen employed in this study. Additionally, the inoculation dose for that study (~10^4^ CFU) was four orders of magnitude lower than our c*Kp* challenge dose. Finally, it is possible our c*Kp* strain KR174 produces more capsule than the strain used in the other study, potentially revealing greater capsule blocking in our experiments. It may be that O-antigen based vaccines are more appropriate for targeting some c*Kp* strains, as increased capsule in hv*Kp* clearly inhibited protection. It remains to be further elucidated if capsule exerts varying degrees of inhibition across a range of c*Kp* strains.

This work provides insight into possible capsule blocking of O-antigen, but it raises several questions that warrant further study. First, we examined single representative K2:O1 c*Kp* and hv*Kp* isolates. Certainly, to gain a deeper understanding of vaccine efficacy and to generalize these findings further would require challenge studies with numerous matched clinical isolates of both pathotypes. Second, there has been some suggestion from previous works that K2 may block O-antigen more effectively than other capsular polysaccharide structures (*21, 22*); therefore, different combinations of K and O types may yield different comparative efficacy results. Third, O-antigen subtypes have recently been described that are structurally and antigenically distinct from each other. For example, O1 O-antigen is comprised of at least two subtypes: O1v1 and O1v2 (*20*). Distinct vaccine candidates need to be developed against both structures to determine efficacy and potential cross-protection. Fourth, our murine models of c*Kp* pneumonia and bacteremia both involved very high doses of KR174, and both vaccines elicited only modest protection in these models. An immunocompromised murine model could be explored to allow for vaccine testing in a system that better mimics the c*Kp* patient population and requires a lower inoculum dose. Further, as capsule is required for lung virulence (*42, 43*) it is not feasible to do a challenge study to test vaccine efficacy with the capsule mutants in lieu of a lower inoculum or immunocompromised model. Fifth, we have recently shown that some c*Kp* infections may not rely solely on antibody responses to fight infection, which could offer an additional explanation for the survival differences we observed in hv*Kp* and c*Kp* survival models (*44*). Finally, the means by which the presence of capsule renders O-antibodies less effective is not mechanistically defined here. Additional studies are needed to specify how capsule blocks O-antibody binding and/or function, including possible effects of altering capsule quantity and structure and potential direct interactions between capsule and O-antigen.

The increasing rates of infection and antibiotic resistance among *K. pneumoniae* represent a serious threat to global health. As such, it is important to define a vaccination strategy to combat this pathogen. Our work here demonstrates that both K- and O-type bioconjugate vaccines are immunogenic and produce functional antibodies, but O-antigen antibodies were less functional in the presence of K2 capsule. Protection from matched capsule-based vaccines may be superior to that of O-conjugate vaccines for a variety of strains, especially hypervirulent strains. Here, we present data supporting the hypothesis of capsule interfering with O-antigen antibody recognition. These data should be corroborated and generalized to additional combinations of *K. pneumoniae* strains, and must be considered when developing a multivalent, efficacious vaccine to target this troublesome pathogen.

## Materials and Methods

### Bacterial strains, plasmids, and growth conditions

Strains and plasmids used in this work are listed in **Table 1**. *K. pneumoniae* strains and mutants were used for all challenge experiments, ELISAs, and serum bactericidal assays. ATCC 43816 was previously identified as a capsule type K2 and O-antigen type O1v1 strain (*45*). The pulmonary clinical isolate KR174 was determined to be capsule type K2 and O-antigen type O1v1 by whole genome sequencing (SeqCenter) and Pathogenwatch (*46*). Primers and oligos used for assembly of DNA constructs in this study are listed in **Supplemental Table 1**. Plasmid pVNM245 expressing the EPA protein containing two sequons and PglS was assembled from a precursor vector pVNM167 expressing EPA with a single sequon and the PglS_ADP1_ gene downstream on a pEXT20 plasmid backbone (*47*). pVNM245 was generated from pVNM167 by separate PCR reactions to amplify products with terminal homology regions: (*i*) the vector backbone with PglS and EPA with one iGT, (*ii*) the second iGT for integration between E548 and G549, and (*iii*) the C-terminus of EPA downstream of the iGT. The DNA fragments were assembled using NEBuilder HiFi DNA Assembly kit (New England Biolabs) and the plasmid was transformed into *E. coli* Stellar (Takara Bio) cells and plated on selective Luria-Bertani (LB) agar with 100 μg/mL ampicillin. Select resulting colonies were grown in LB with ampicillin overnight, plasmids extracted using a GeneJet Plasmid Purification Kit (Thermo Scientific), and constructs sequence-verified using Sanger sequencing (Genewiz).

**Table 1.**
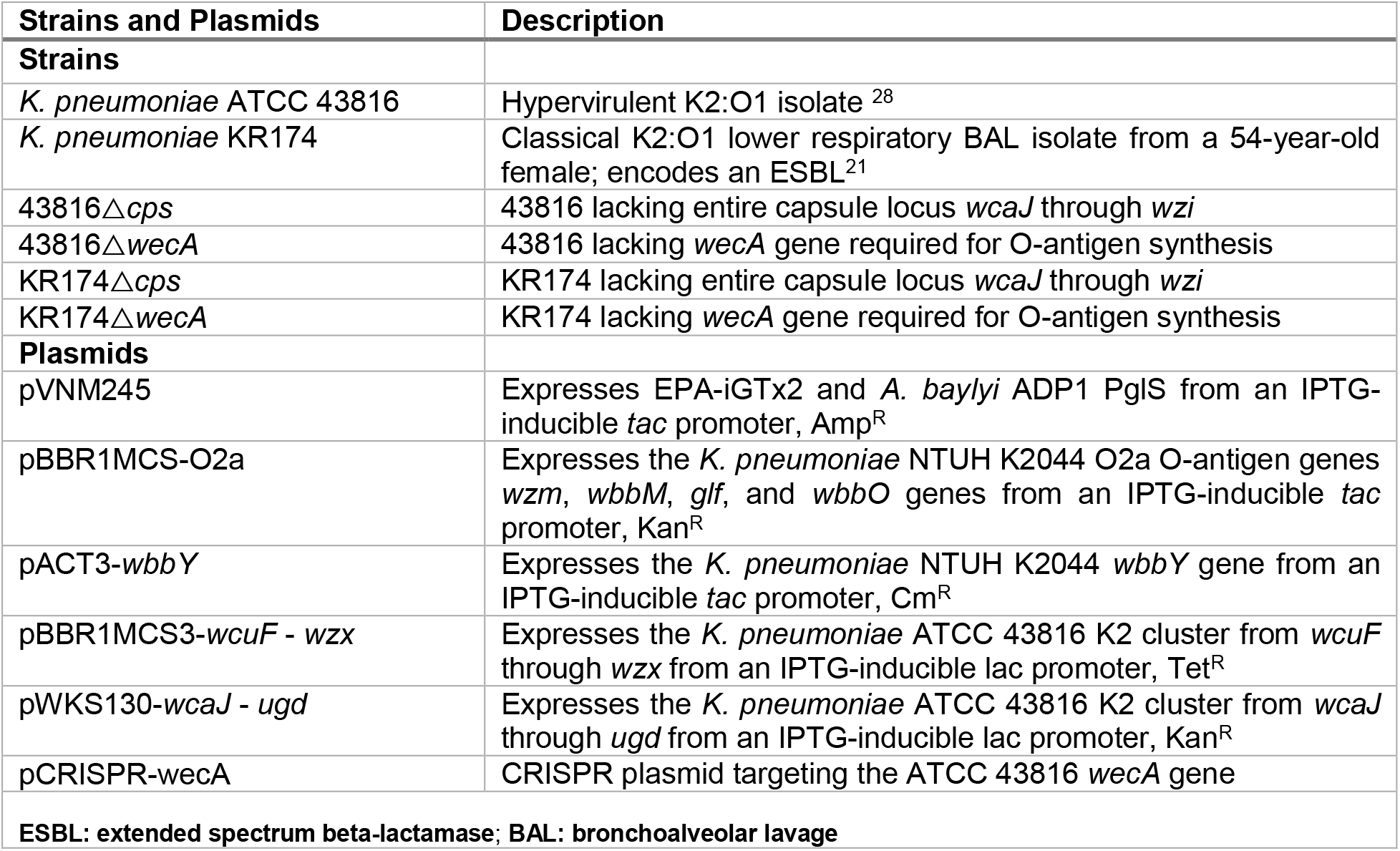
Strains and plasmids used in this study.

### Construction of K2 and O1 polysaccharide expression plasmids

The *K. pneumoniae* K2 capsule cluster from ATCC 43816 was cloned in two separate sections using In-Fusion Snap Assembly (Takara Bio USA). The genes from *wcuF* through *wzx* were cloned into PCR-linearized pBBR1MCS3 (*48*), and the genes from *wcaJ* through *ugd* were cloned into KpnI-digested pWKS130 (*49*). The genes required for producing *K. pneumoniae* O1 O-antigen (*24*) in *E. coli* were cloned into two separate expression vectors. The *wzm, wbbM, glf, wbbO*, and *wbbY* genes were amplified from *K. pneumoniae* NTUH K2044 genomic DNA with complementary 20 – 30 bp overhangs for Gibson assembly with PCR-linearized pBBR1MCS2 (*48*) for *wzm* – *wbbO* or XbaI restriction-digested pACT3 (*47*) for *wbbY*.

### Production of bioconjugates in E. coli

To produce O1 bioconjugates, the pBBR1MCS2-O2a and pACT3-*wbbY* plasmids were electroporated into CLM24 *E. coli* (*27*) harboring pVNM245. K2 bioconjugates were produced by electroporating K2 expression vectors into SDB1 *E. coli* (*27*) hosting pVNM245. The cells were outgrown for 1 h at 37 °C and then plated on LB agar with appropriate antibiotics (100 μg/mL ampicillin, 20 μg/mL kanamycin, and/or 20 μg/mL chloramphenicol). The following day, 8 – 10 resulting colonies were inoculated into LB, incubated overnight at 30 °C, and next morning subcultured into 2 L flasks with 1 L Terrific Broth (TB) media. Cells were cultured while shaking for 3-4 h to reach OD_600_ of 0.4-0.5, then induced with 0.1 mM isopropyl β-D-1-thiogalactopyranoside (IPTG) and grown overnight. Cells producing O1 and K2 bioconjugates were grown at 25 °C and 30 °C after induction, respectively. After 24 h total growth, cells were harvested and prepared for protein purification.

### Periplasmic extraction and purification of EPA, EPA-O1 and EPA-K2

Unglycosylated carrier protein and bioconjugates were purified from *E. coli* periplasmic extracts. *E. coli* frozen cell pellets were thawed in 0.9% NaCl saline solution and centrifuged at 6,000 x *g* for 20 min. Each washed pellet was resuspended in hyperosmotic buffer containing 200 mM Tris-HCl pH 8.5, 100 mM EDTA, and 25% w/v sucrose and gently rolled for 45 min at 4 °C. Cells were centrifuged at 10,000 x *g* for 20 min and the recovered pellet was suspended in hypoosmotic buffer containing 20 mM Tris-HCl pH 8.5 and gently rolled for 45 min at 4 °C. The periplasmic extract was recovered by spinning the suspension at 18,000 x *g* for 30 min. The resulting supernatant was filtered through a 0.45-μm PES membrane and loaded onto pre-equilibrated Nickel NTA Agarose Beads (Goldbio). The beads were washed with equilibration/wash buffer containing 20 mM Tris-HCl pH 8, 500 mM NaCl, and 10 mM imidazole. The protein was then eluted in the same buffer containing 300 mM imidazole and buffer-exchanged into 20 mM Tris-HCl pH 8 using a 50-kDa cutoff PES centrifugal concentrator (Pierce). The protein was loaded onto a SourceQ 15Q 4.6/100 PE anion-exchange column using an Äkta pure 25L FPLC instrument (GE Healthcare). For all conjugates, Buffer A contained 20 mM Tris-HCl pH 8.0 and Buffer B contained 20 mM Tris-HCl pH 8.0 and 1 M NaCl. The proteins were eluted using a step gradient running at a constant 2 mL/min. O1-EPA was eluted using a step gradient from 0 – 5 – 7.5 – 20% B with 8 column volumes per step. O1-EPA eluted in the 5 and 7.5% fractions. Unglycosylated EPA and K2-EPA were purified from separate purifications using the same 10 – 15 – 20 – 25% B step gradient with 8 column volumes per step. K2-EPA eluted in the 20 and 25% B fractions and unglycosylated in the 10 and 15% B fractions. The pooled fractions from anion-exchange were concentrated and loaded on an S200 Increase 10/300 GL size-exclusion column in 1X PBS buffer running at a constant flow rate of 0.75 mL/min. The purified final protein concentration was determined using a Pierce BCA Protein Assay Kit.

### Nuclear magnetic resonance (NMR) spectroscopy

K2 and O1 polysaccharides heterologously expressed in glycoengineered *E. coli* were extracted by heating whole-cell bacteria in 2% acetic acid at 105 °C for 1.5 h. Insoluble material was removed by centrifugation, and the supernatant (containing K2 or O1 polysaccharide) was isolated on Sephadex G50 column, subsequently purified on Hitrap Q (acidic) column, polished on a Hitrap S column to remove trace protein background (for the O1 polysaccharide) or desalted on a Sephadex G15 column (for the K2 polysaccharide). Polysaccharides were dried and analyzed by NMR. NMR experiments were carried out on a Bruker AVANCE III 600 MHz (^1^H) spectrometer with 5 mm Z-gradient probe with acetone internal reference (2.23 ppm for ^1^H and 31.45 ppm for ^13^C) using standard pulse sequences cosygpprqf (gCOSY), mlevphpr (TOCSY, mixing time 120 ms), roesyphpr (ROESY, mixing time 500 ms), hsqcedetgp (HSQC), hsqcetgpml (HSQC-TOCSY, 80 ms TOCSY delay) and hmbcgplpndqf (HMBC, 100 ms long range transfer delay). Resolution was kept <3 Hz/pt in F2 in proton-proton correlations and <5 Hz/pt in F2 of H-C correlations. The spectra were processed and analyzed using the Bruker Topspin 2.1 program. Monosaccharides were identified by COSY, TOCSY, and NOESY cross peak patterns and ^13^C NMR chemical shifts. Connections between monosaccharides were determined from transglycosidic NOE and HMBC correlations.

### Mass spectrometry

Intact mass analysis was performed on a Xevo G2-XS QTof Quadrupole Time-of-Flight Mass Spectrometer coupled to an ACQUITY UPLC H-class system (Waters) using a Jupiter 300 C5 column (2mm*50mm, Phenomenex). Protein samples were resuspended in 2% acetonitrile, 0.1% trifluoroacetic acid and loaded/separated on the C5 column at a flow rate of 0.25 mL/min. 2 μg of each bioconjugate (K2-EPA and O1-EPA) was desalted on column for 2 min with Buffer A (2% acetonitrile 0.1% formic acid) before being separated by altering the percentage of Buffer B (80% acetonitrile, 0.1% formic acid) from 0% to 100% over 10 min. The column was then held at 100% Buffer B for 0.5 min before being equilibrated for 1 min with Buffer A, for a total run time of 13.5 min. Samples were infused into the Xevo G2-XS QTof Quadrupole Time-of-Flight Mass Spectrometer using electrospray ionization (ESI) and MS1 mass spectra acquired with a mass range of 400–2000m/z at 1 Hz. Scans across the apex of the elution peaks were summed then peak lists exported. Deconvolution of proteoforms within K2-EPA and O1-EPA were undertaken using UniDec (*50*).

### Construction of K. pneumoniae mutant strains

A modified Red recombinase protocol was utilized to construct the Δ*cps* mutants (lacking the *cps* operon from *wzi* to *wcaJ*) using pKD46s (*28*). Linear regions of DNA were amplified from pKD4 using primers listed in **Supplemental Table 1**, resulting in the region of interest being replaced with a kanamycin cassette. All mutants were confirmed by sequencing of amplicons generated via PCR using check primers. Δ*wecA* mutants were constructed using a modified CRISPR method (*51*). The zeocin resistance cassette of pUC19_CRISPR_DpmrA was replaced with a tetracycline resistance gene, and the *pmrA* gRNA and homology arms were swapped for those targeting *wecA*. Deletion mutants lacking *wecA* were identified by PCR and confirmed by sequencing.

### Glucuronic acid quantification

Quantification of capsule was performed using glucuronic acid assays, as previously described (*52*). Briefly, bacteria were grown overnight in LB broth at 37 °C under static growth conditions. Cultures were pelleted and resuspended in 1x PBS to an OD_600_ of 1.0. A total of 500 μL of normalized culture was mixed with 100 μL of 1% Zwittergent 3-14 (Sigma) in triplicate in 100 mM citric acid and incubated at 50 °C for 20 min. After centrifugation, supernatants from samples were precipitated with cold ethanol at 4 °C for 20 min. Upon precipitation, samples were recentrifuged and the pellets were dissolved in 200 μL sterile water and 1200 μL of 12.5 mM tetraborate in concentrated H_2_SO_4_. Samples were vortexed, boiled at 95 °C for 5 min, and mixed with 20 μL of 0.15% 3-hydroxydiphenol (Sigma) in 0.5% NaOH. Absorbance was measured at 520 nm using a microplate reader (Bio-Tek). The uronic acid concentration of each sample was determined using a standard curve of glucuronic acid (Sigma). The limit of detection (LOD) was previously defined (*53*) and reported here. Significance was determined using Mann-Whitney nonparametric tests with p<0.05. All graphs and statistics were generated using GraphPad Prism version 9.

### Centrifugation assay

Levels of hypermucoviscosity were determined as previously described (*28*). Bacteria were cultured standing overnight in LB media at 37 °C. Cultures were pelleted and resuspended in PBS to an OD_600_ of 1.0. One mL of normalized culture was centrifuged at 500 *x g* for 5 min, and OD_600_ of the resulting supernatant was measured. Final readings were normalized to the starting OD_600_ value. Significance was determined using Mann-Whitney nonparametric tests with p<0.05. All graphs and statistics were generated using GraphPad Prism version 9.

### Murine vaccination

All murine immunizations complied with ethical regulations for animal testing and research. Experiments were carried out at Washington University School of Medicine in St. Louis according to the institutional guidelines and received approval from the Institutional Animal Care and Use Committee at Washington University in St. Louis. Five-week-old female BALB/c mice (Jackson Laboratories) were subcutaneously injected with 100 μL of a vaccine formulation on days 0, 14, and 28. The vaccination groups were as follows: EPA carrier protein alone, K2-EPA and O1-EPA. All vaccines were formulated with Aldydrogel® 2% aluminum hydroxide gel (InvivoGen) at a 1:9 ratio (50 μL vaccine to 5.5 μL adjuvant in 44.5 μL sterile PBS). All vaccination groups received 1 μg of vaccine based on total polysaccharide content. The total polysaccharide content was measure using a modified anthrone-sulfuric assay (*54*). Sera were collected on days 0, 14, 28, and 42 prior to immunizations or challenge. Mice were challenged with either hv*Kp* or c*Kp* on day 42 (described below).

### Enzyme-linked immunosorbent assays

Briefly, 96 well plates (BRAND™ immunoGrade microplates) were coated overnight with ~10^6^ CFU/100 μL of the specified *K. pneumoniae* strain in sodium carbonate buffer. All strains and growth conditions are described above. After coating, wells were blocked with 1% BSA in sterile PBS and washed with 0.05% PBS-Tween-20 (PBS-T); all subsequent washes were the same. Sera from immunized mice were diluted 1:100 and added to wells in triplicate for 1 h at room temperature. After washing, HRP-conjugated anti-mouse IgG (GE Lifesciences, 1:5000 dilution in PBS-T) was added to wells for 1 h at room temperature. Plates were washed, developed using 3,3’,5,5’-tetramethyl benzidine (TMB) substrate (Biolegend), and stopped with 2 N H_2_SO_4_. Absorbance was determined at 450 nm using a microplate reader (Bio-Tek). Total IgG concentration was determined using an IgG standard curve. Standard wells were coated with diluted mouse IgG in sodium carbonate buffer and treated the same as sample wells thereafter. All wells were normalized to blank wells coated and treated the same as sample wells without receiving primary mouse sera. Significance was determined using Mann-Whitney nonparametric tests with p<0.05. All graphs and statistics were generated using GraphPad Prism version 9.

### Western blots

OD_600_-normalized whole-cell lysates of wild-type strains (43816 and KR174) and subsequent mutants (Δ*cps* and Δ*wecA*) were separated on 7.5% Mini-PROTEAN precast polyacrylamide gels (Biorad). Samples were transferred to nitrocellulose membranes (Bio-Rad) and blocked with LI-COR Intercept blocking buffer for 45 min. Membranes were incubated with primary antibody for 30 min, washed three times with TBS 0.1% v/v Tween-20 (TBST), incubated with secondary antibody for 30 min, washed three times with TBST, and visualized using an Odyssey Infrared Imaging System (LI-COR Biosciences). Primary antibodies were pooled day 42 mouse sera from mice immunized with either K2-EPA bioconjugate or O1-EPA bioconjugate used at 1:1000 concentration. Secondary antibody was LI-COR IRDye 680RD goat anti-mouse (925-68070) used at 1:10,000 dilution. For western blots of purified proteins, 0.75 μg protein was loaded per lane and prepared as above. Primary antibodies used at 1:1,000 dilution were commercial rabbit anti-EPA antibodies (Millipore-Sigma), and O1 antibodies (*20*) were a generous gift from Prof. Chris Whitfield (University of Guelph). Secondary antibodies for purified proteins were LI-COR IRDye 680RD goat anti-mouse (925-68070) and Licor 800CW goat anti-rabbit (925-32211) both at 1:10,000 dilution. For western blots of LPS, 15 μL of the LPS mini-prep was separated on a 4-20% TGX gel (Bio-Rad) and prepared as above. Primary antibody was day 42 mouse sera from mice immunized with the K2-EPA bioconjugate used at 1:1,000 dilution. The secondary antibody for LPS western blotting was LI-COR IRDye 680RD goat anti-mouse (925-68070) at 1:10,000 dilution.

### LPS mini-prep extraction and silver staining

LPS was extracted, as previously described (*55*). Briefly, LPS was extracted using a hot phenol method from 2.0 OD units of glycoengineered *E. coli*. LPS was resuspended in 50 μL of Laemmli buffer. A 15 μL aliquot of LPS was separated on a 15% sodium dodecasulfate (SDS) polyacrylamide gel. Silver staining of SDS gels was performed, as previously described (*56*).

### Serum bactericidal assays

Serum bactericidal assays for *K. pneumoniae* were adapted from methods previously described (*57*). *K. pneumoniae* isolates and derived mutants were grown under static conditions in LB broth for 16 hours at 37°C. Cultures were centrifuged at 8000 *x g* for 10 min and resulting pellets were resuspended in sterile PBS to an OD_600_ ~1.0. Cultures were diluted 1:80,000 in sterile PBS. The assay mixture was prepared in a 96-well U-bottom microtiter plate (TPP®) by combining 70 μL of diluted bacteria and 20 μL of diluted heat-inactivated mouse serum. Sera were day 42 from mice immunized with EPA, K2-EPA, or O1-EPA bioconjugates and heat inactivated at 56 °C for 30 min. After incubation at 37 °C with shaking for 1 h, 10 μL of baby rabbit complement (Pel-Freez Biologicals) was added to wells at a final concentration of 10% and incubated for an additional 1 h at 37 °C with shaking. Control wells were treated the same as samples except for receiving diluted heat-inactivated pre-immune mouse serum. After the final incubation, samples were serially titrated in sterile PBS and plated in pentaplicate. Colonies were counted after 16-h incubation at room temperature. SBA titers were calculated as surviving colonies compared to the control multiplied by 100 to determine percent survival. Statistical analyses were performed using Mann-Whitney test with p<0.05. All graphs and statistics were generated using GraphPad Prism version 9.

### Murine challenge experiments

For the pneumonia model, groups of vaccinated mice were anesthetized with isoflurane and inoculated with bacterial suspension via oropharyngeal aspiration as previously described (*8, 58*). For the bacteremia model, isolates were administered via intraperitoneal injection. Strain 43816 was grown statically in LB broth for 16 h at 37 °C. Cultures were centrifuged at 8000 *x g* for 10 min, and pellets were resuspended in sterile PBS to an OD_600_ ~1.0. Cultures were diluted 1:80,000 in sterile PBS to obtain the desired final concentration. Challenge doses of 43816 were ~1500 CFU in 50 μL for aspiration and ~2000 CFU in 50 μL for bacteremia, respectively. KR174 was grown statically in LB broth for 16 h at 37 °C. Cultures were centrifuged at 8000 *x g* for 10 min, and pellets were resuspended in sterile PBS to an OD_600_ ~2.0 (for aspiration inoculum) or ~1.0 (for bacteremia inoculum). Challenge doses of KR174 were ~10^9^ CFU in 50 μL for aspiration and ~10^8^ CFU in 50 μL for bacteremia, respectively. Mouse survival and weight were monitored daily for two weeks. Each experiment was performed in duplicate with n=10 mice per group. Pairwise survival differences were determined by the Log-rank (Mantel-Cox) test. All graphs and statistics were generated using GraphPad Prism version 9.

### Blood titers

To quantify bacteremia, female BALB/c mice (5 mice per group) were injected i.p. with 50 μL of either 43816 or KR174, as described above. 24 hours after injection, mice were euthanized and blood samples were taken via cardiac puncture. Samples were serially titrated in sterile PBS and plated in pentaplicate. Colonies were counted after 16-h incubation at room temperature.

## Acknowledgments

We would like to thank Dr. David Hunstad for critical review of this manuscript. This work was supported by the Children’s Discovery Institute of Washington University and St. Louis Children’s Hospital (to DAR). PLW was supported by the W.M. Keck Fellowship. This work was also supported by the US National Institute of Allergy and Infectious Diseases (R41 AI167078-01 and R42 AI165116-01A1 to CMH). NES is supported by an Australian Research Council Future Fellowship (FT200100270) and an ARC Discovery Project Grant (DP210100362). We thank the Melbourne Mass Spectrometry and Proteomics Facility of The Bio21 Molecular Science and Biotechnology Institute for access to MS instrumentation.

---

**Fig. S1.**
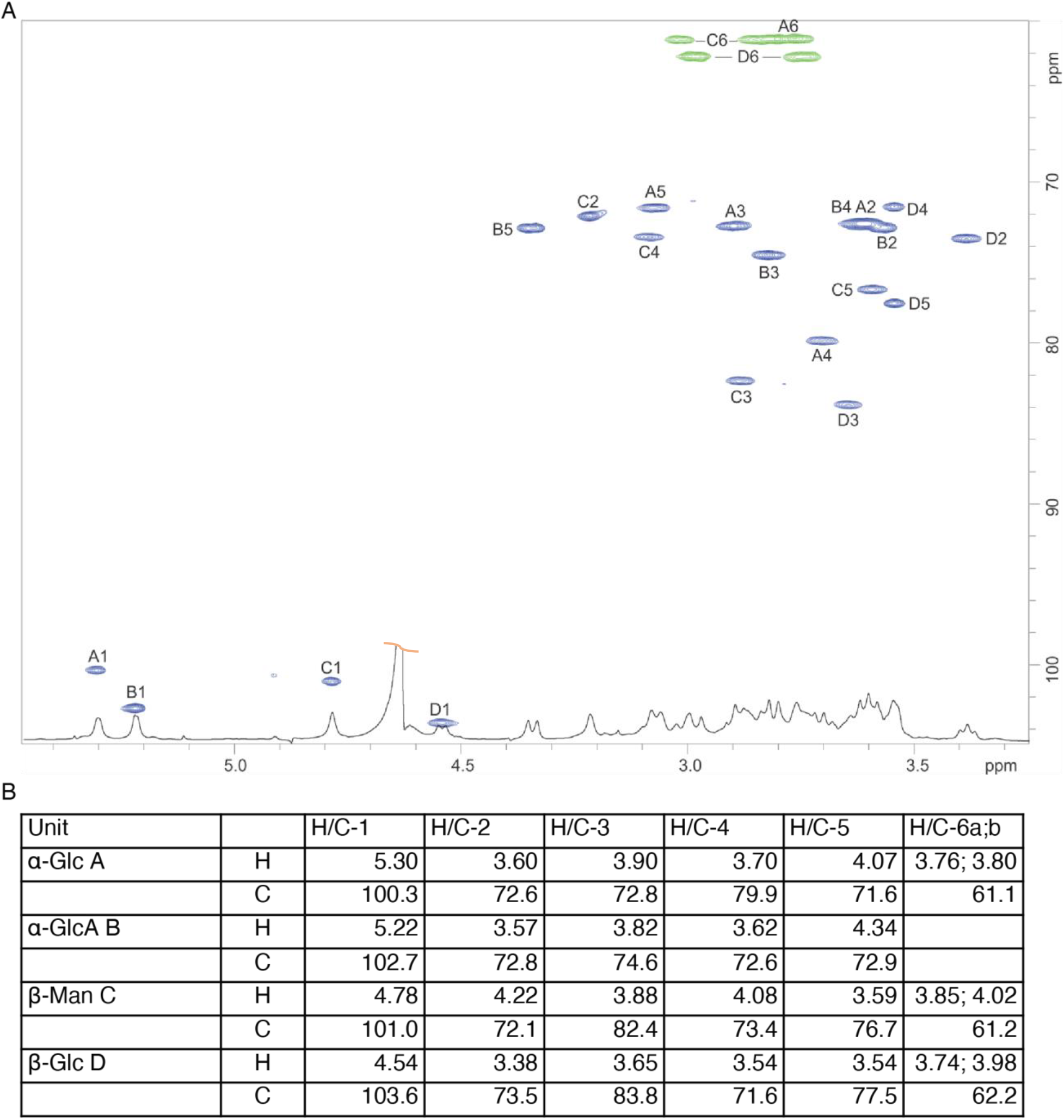
Two-dimensional NMR analysis of K2 polysaccharide extracted from glycoengineered *E. coli* expressing the decoupled K2 gene cluster. (A) ^1^H-^13^C HSQC spectrum of the extracted K2 polysaccharide. Orange hash indicates cut water peak. (B) NMR data for the K2 polysaccharide (D_2_O, 25 °C, 600 MHz).

**Fig. S2.**
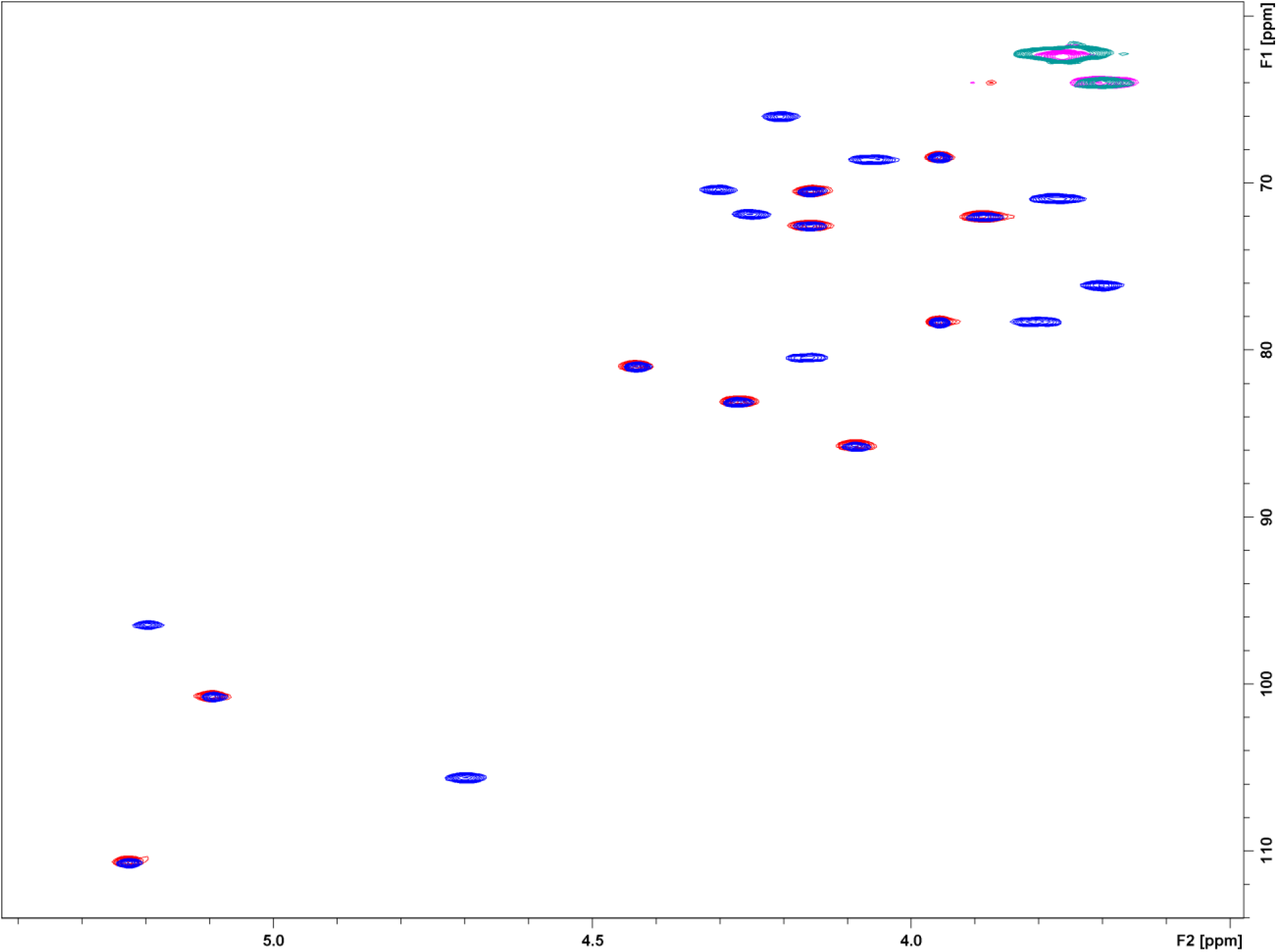
Two-dimensional NMR spectroscopy of the O1 polysaccharide heterologously expressed in glycoengineered *E. coli*. Overlap of the ^1^H-^13^C HSQC spectra of *Klebsiella* O1, galactan II constituent (blue-cyan), and galactan I constituent (red-pink), polysaccharide portions.

**Fig. S3.**
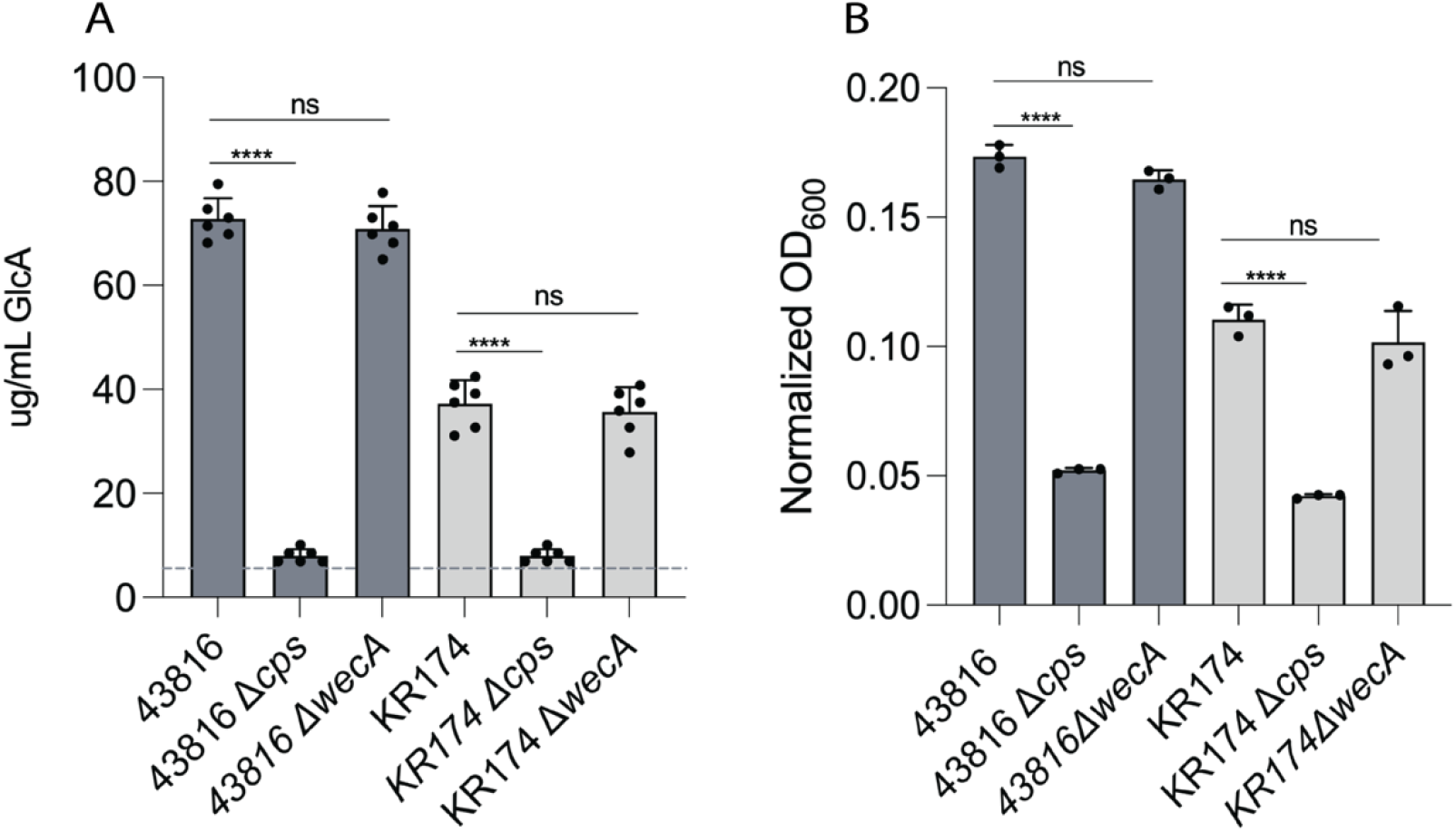
Classical and hypervirulent *K. pneumoniae* capsule and hypermucoviscosity phenotypes. (A) Capsule quantification using a glucuronic acid assay comparing hv*Kp* 43816), c*Kp* KR174, and their respective capsule (*cps*) and O antigen (*wecA*) knockouts. The dotted line represents the limit of detection for uronic acid. (B) Hypermucoviscosity quantification via low-speed centrifugation assay. Statistical analyses were performed via Mann-Whitney U test. **** p<0.0001; ns, not significant.

**Fig. S4.**
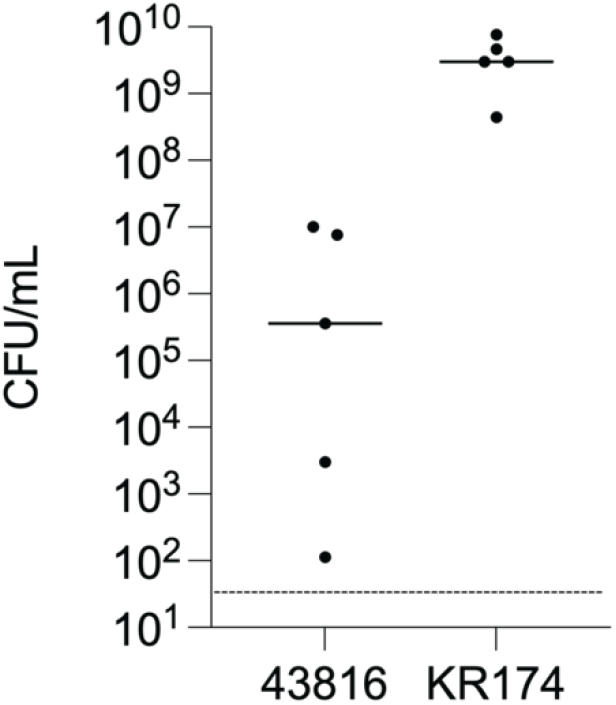
Bacteremia resulting from intraperitoneal challenge with 43816 or KR174. BALB/c mice were infected with 2000 CFU 43816 or 10^8^ CFU KR174 in 50 μL via intraperitoneal injection, and blood was collected for culture after 24 h. The dotted line represents the limit of detection.

**Supplemental Table 1:**
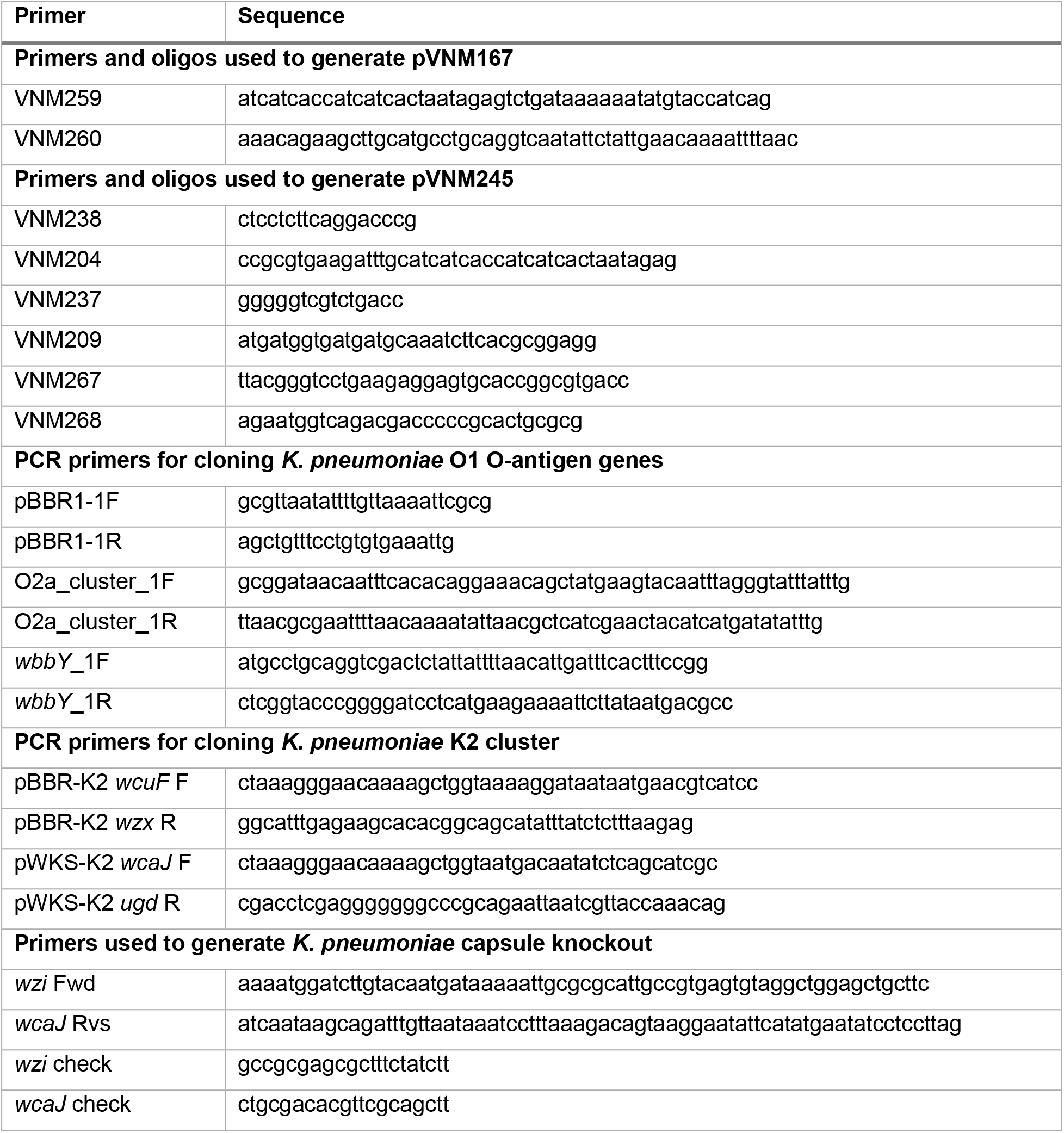
Primers and oligos used in this study.

